# Living to the high extreme: unraveling the composition, structure, and functional insights of bacterial communities thriving in the arsenic-rich Salar de Huasco – Altiplanic ecosystem

**DOI:** 10.1101/2021.04.13.439755

**Authors:** J Castro-Severyn, C Pardo-Esté, KN Mendez, J Fortt, S Marquez, F Molina, E Castro-Nallar, F Remonsellez, CP Saavedra

## Abstract

Microbial communities inhabiting extreme environments like Salar de Huasco (SH) are adapted to thrive while exposed to several abiotic pressures and the presence of toxic elements like arsenic (As). Hence, we aimed to uncover the role of arsenic in shaping bacterial composition, structure, and functional potential in five different sites in this Altiplanic wetland using a shotgun metagenomic approach. The sites exhibit wide gradients of arsenic (9 to 321 mg/kg), and our results showed highly diverse communities and a clear dominance exerted by the *Proteobacteria* and *Bacteroidetes* phyla. Functional potential analyses showed broadly convergent patterns, contrasting with their great taxonomic variability. Arsenic-related metabolism is different among the five communities, as well as other functional categories like those related to the CH_4_ and S cycles. Particularly, we found that the distribution and abundance of As-related genes increase, following along the As concentration gradient. Approximately 75% of the detected genes for As-metabolism belong to expulsion mechanisms, being *arsJ* and *arsP* pumps related to sites with higher As concentrations and present almost exclusively in *Proteobacteria*. Furthermore, taxonomic diversity and functional potential are reflected in the 12 reconstructed high-quality MAGs (Metagenome Assembled Genomes) belonging to *the Bacteroidetes* (5), *Proteobacteria* (5), *Cyanobacteria* (1) and *Gemmatimonadota* (1) phyla. We conclude that SH microbial communities are diverse and possess a broad genetic repertoire to thrive under extreme conditions, including increasing concentrations of the highly toxic As. Finally, this environment represents a reservoir of unknown and undescribed microorganisms, with a great metabolic versatility, which needs further study.

**IMPORTANCE:** Microbial communities inhabiting extreme environments are fundamental for maintaining the ecosystems; however, little is known about their potential functions and interactions among them. We sampled the microbial communities in Salar de Huasco (SH) in the Chilean Altiplano, a fragile and complex environment that comprises several stresses. We found that microbes in SH are taxonomically diverse; nonetheless, their functional potential seems to have an important convergence degree, suggesting high adaptation levels. Particularly, arsenic metabolism showed differences associated with increasing concentrations of the metalloid throughout the area, and it is effectively exerting a clear and significant pressure over these organisms. Thus, this research’s significance is that we described highly specialized communities thriving in little-explored environments under several pressures, considered analogous of early Earth and other planets, and can have the potential for unraveling technologies to face climate change repercussions in many areas of interest.

## INTRODUCTION

Extreme environments like high-altitude wetlands select for adaptations in bacterial communities that enable them to thrive. This particular and fragile environment resemble life before the oxygenation of Earth and could serve as models for studying life in other planets. Moreover, microbial communities are critical for maintaining biogeochemical cycles, particularly in extreme environments where there is little presence of other life forms. Therefore, microorganisms are crucial for ecosystems’ health and functioning (1). Salar de Huasco (SH), a high-altitude wetland located in the Chilean Altiplano (20°18’18’’S; 68°50’22’’W, Chile) at 3,800 m.a.s.l. is a Ramsar protected site, considered a hotspot for microbial life (2, 3). This area is labeled as extreme due to very particular confluence of physicochemical and environmental conditions, like negative water balance, high daily temperature variation, very arid conditions, high salinity, low atmospheric pressure, high solar radiation, presence of arsenic, among other stressors (4, 5, 6, 7, 8, 9).

In high altitude wetlands, arsenic concentration, moisture availability, and salt concentrations model communities at a small scale (10, 11, 12, 13). The presence of toxic metal(oids), such as arsenic, is one of the main drivers of microbial communities’ composition (14), as an important selection pressure originated from natural geochemical processes or human activities. Also, previous studies have determined that the presence of metal(oids) can influence biogeochemical cycles, namely C, N and S, by promoting specific chemical reactions and the enrichment of chemoautotrophs, for example As(III)-oxidizing bacteria that can couple this process with nitrate reduction (15, 16, 17); thus, making relevant to assess the metabolic potential of indigenous microbes and communities. In the North of Chile, the relationship between volcanism activities and the presence of arsenic is a known feature, this geological process has been attributed to hydrothermal conditions such as geysers and fumaroles in the pre-range and high plateau (5). Besides, bacteria that thrive in these environments exhibit arsenic-related genes, including genes associated with methylation (ArsM), oxidation (AioAB, ArxAB), and dissimilatory reduction (ArrAB) (18). The most common resistance mechanism is based on As(III) extrusion from the cell by efflux pumps, which is commonly coupled with As(V) reduction (19). Thus, making the genes from the *ars* operon the most abundant arsenic resistance markers; the basic requirements comprises the arsenite efflux (ArsB, Acr3) and arsenate reductase (ArsC) (20).

Arsenic is a crucial element for the microbial community composition and metagenomic approaches have helped to shed some light on this subject. For instance, in Socompa’s stromatolites arsenic resistance is achieved mainly through reduction and expulsion of As(V) via Acr3 efflux pumps reductases (21, 22). Additionally, others have found that the presence of arsenic influenced the microbial communities, and the *Rhodococcus* genus was significantly enriched in elevated levels of the metalloid in groundwater (14). Furthermore, arsenic mobilization is distributed across a broad phylogenetic lineage in similar ecosystems, as *arrA* was detected in *Betaproteobacteria, Deltaproteobacteria*, and *Nitrospirae* MAGs (23). Also, although arsenic-related genes are widespread, they are not universal, in particular As metabolism genes such as *aioA, arrA, arsM, arxA* are less common in the environment compared with *acr3, arsB, arsC* and *arsD* (24).

In this study, we tested different established sites in the SH regarding taxonomic and functional heterogeneity. We focused on As-related genetic elements as well as other relevant metabolic functions. Evermore, such analyses enable the identification of relatively small subsets of markers associated with a particular ecologically important function. Besides, identifying those and the changes of bacterial communities are potentially useful bio-indicator for monitoring ecosystem health (25, 26). Therefore, considering the aforementioned we set up to describe and characterize the composition, structure and functional potential of bacterial communities from the sediments of five different Salar de Huasco sites along an arsenic gradient, among other environmental pressures.

## RESULTS

The Salar de Huasco comprises an important level of variation (27, 28), evidenced within a relatively small area (sampled area spans a distance of 5.9 km); including daily oscillations in a wide range of environmental parameters (temperature and humidity) and other which vary in spatial gradients, i.e., arsenic and salinity increasing from north to south (Figure 1) (9). In this context, shotgun metagenomic sequencing of the sediment samples representing the five SH sites yielded an average of 87.2 million reads (150 bp length) per sample, with a quality score ≥ 30 presented by ∼95% of those obtained reads.

**Figure 1.**
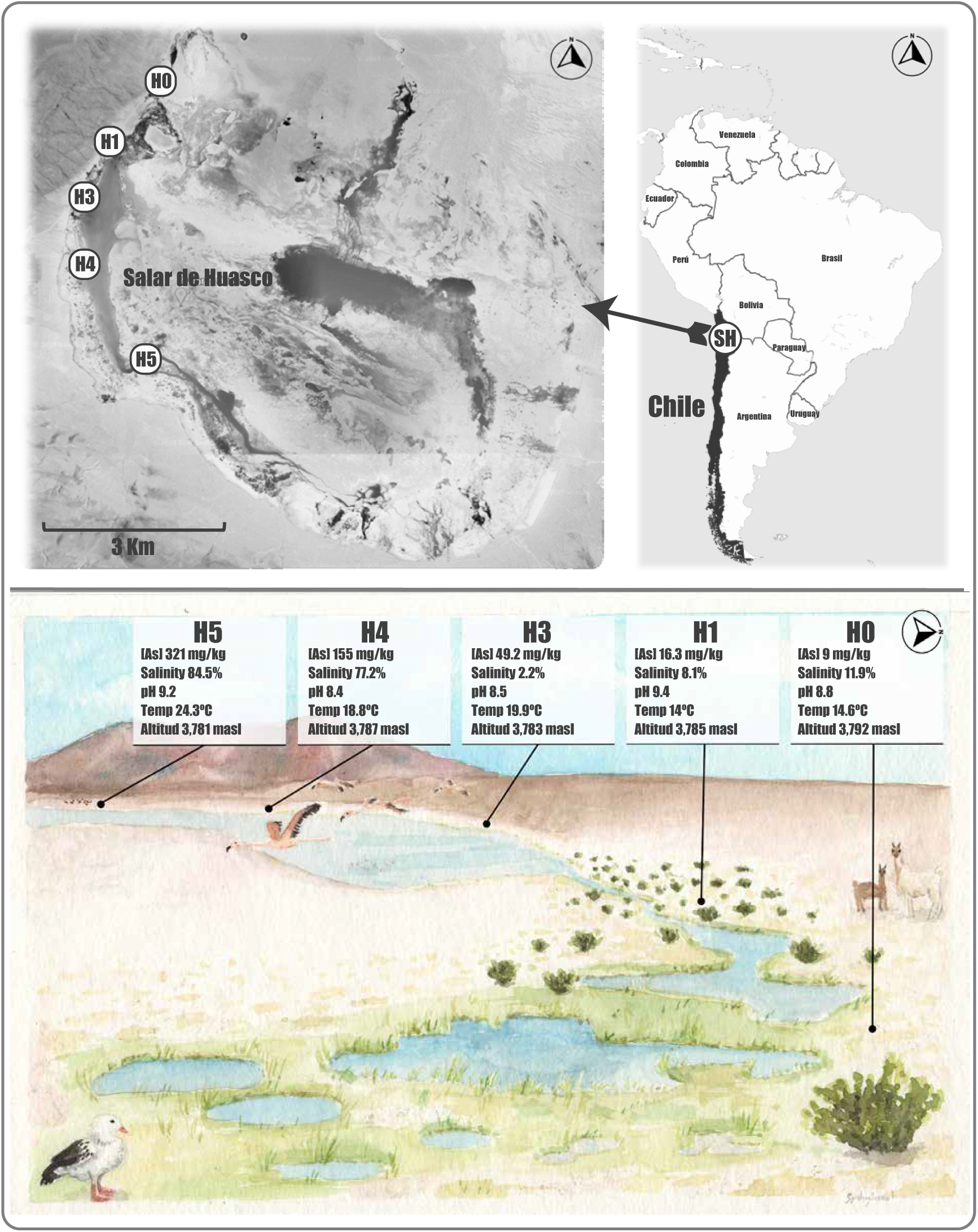
Salar de Huasco study area. Top panel: a map showing the five sampling sites (H0: 20°15′48.8″S - 68°52′28.4″W; H1: 20°16′27.7″S - 68°53′3″W; H3: 20°16′59.2″S - 68°53′16.7″W; H4: 20°17′40.9″S - 68°53′17.3″W and H5: 20°18′37″S - 68°52′42″W) investigated in this study. SH is located between the 68°47′ - 68°54′ W and 20°15′ - 20°20′ S in the Tarapacá region Northern Chile (Source: Google Earth). Bottom panel: illustration of the SH landscape seen from the H0 site to the South-West; the signs show some particular characteristics of each site (according to those reported in Castro-Severyn et al., 2020 (9)). The painting is the work of the illustrator Florence Gutzwiller @spideryscrawl.

### The bacterial communities of SH are highly heterogeneous and rich in unknown taxa

The contrasting differences presented among the SH sampling sites are reflected by the taxonomic composition of bacterial communities, which showed contrasting patterns and variation between sites (Figure 2). Overall, our results indicate that the *Proteobacteria* and *Bacteroidetes* are the most prevalent phyla in the sampled sediments, which are particularly enriched in the H3, H4 and H5 communities, accounting together for >60% of all observed taxa (Figure 2A). In turn, these phyla represent ∼50% in the H0 and H1 communities. Interestingly, the H0 site is dominated by the *Cyanobacteria* phylum with a 34% of the total community; also, this community has the highest abundance of *Firmicutes, Patescibacteria* and *Spirochaetes* phyla as well as significantly lower proportion of *Actinobacteria.* Moreover, H1 community profile presents the highest abundance of *Chloroflexi, Actinobacteria, Verrucomicrobia, Planctomycetes* and *Acidobacteria* phyla. On the other hand, H3, H4 and H5 communities are more similar to each other, despite the vast differences in low-abundance taxa; many of which are exclusively present in only one community (Supplementary Table S1).

**Figure 2.**
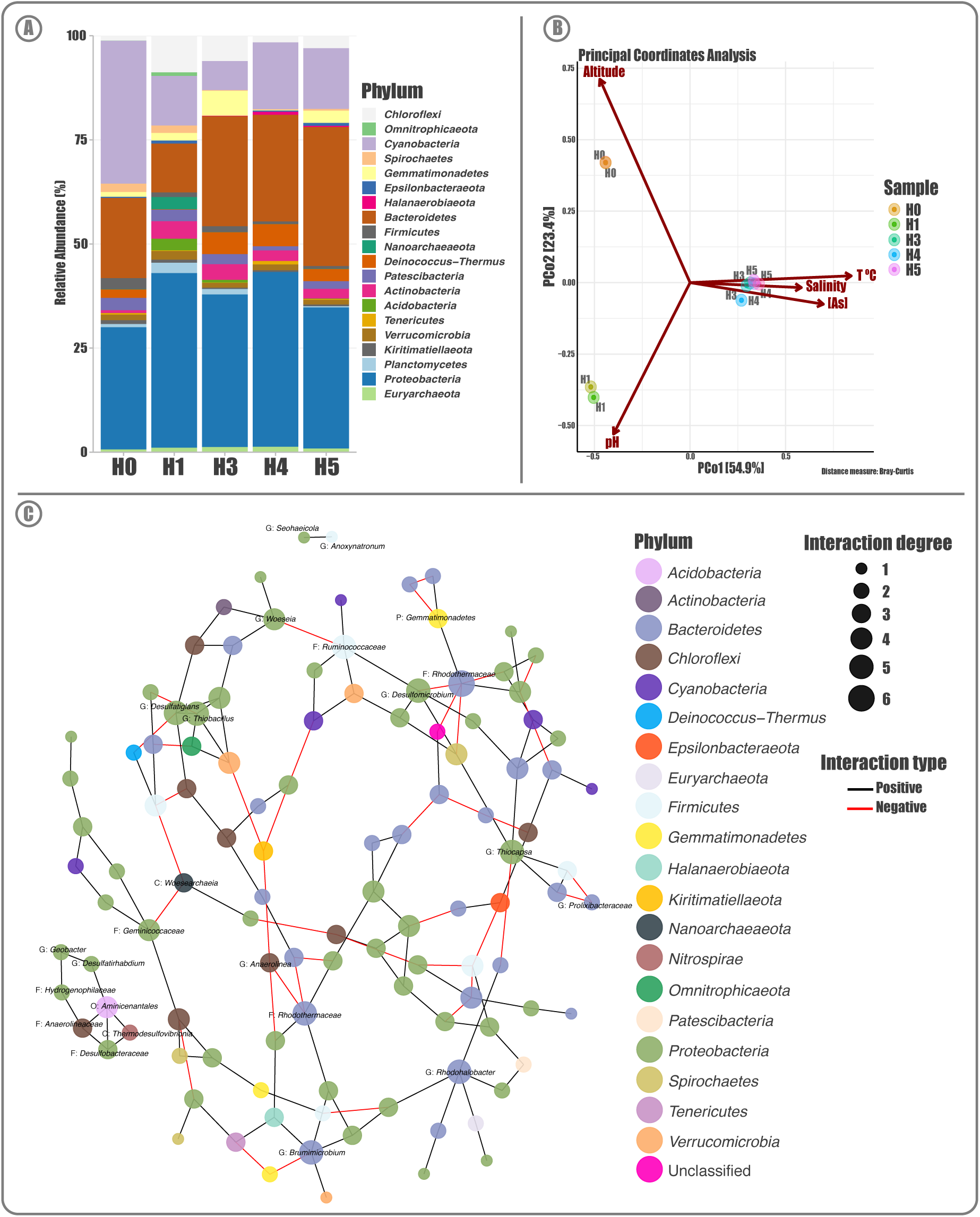
Salar de Huasco bacterial communities. **A)** Taxonomic composition and relative abundance of SH microbial communities in the five studied sites; Stacked-bars show the top20 most abundant bacteria at the phylum taxonomic level. **B)** Beta diversity by Principal Component Analysis (PCoA) on Hellinger transformed amplicon sequence variant (ASV) relative abundances. Each point corresponds to a community from the different sites (represented colors), and its relative distance indicates the level of similarity to all other samples. The arrows indicate the explanatory power of the statistically significant environmental parameters with regards to the observed variation in community composition. **C)** Co-occurrence network analysis of the SH bacterial communities, the size of each node (representing ASVs) is proportional to the number of connections (degrees), the color of the edges connecting nodes represent the interaction type, the node color indicates the taxonomic affiliation at phylum level and node labels are at the lowest available taxonomic classification.

The most abundant bacteria at the lowest (available) taxonomic rank reflect the same pattern, with a great abundance of *Proteobacteria* genera *Roseovarius* and *Desulfotignum* in the H3, H4 and H5 communities (Supplementary Figure S1). Moreover, *Halomonas, Thiobacillus, Luteolibacter* and *Truepera* genera are more widespread, while *Rhodohalobacter, Marinobacter, Psychroflexus* and *Brumimicrobium* are also concentrated in H3, H4 and H5. Contrary, H0 and H1 are enriched in *Cyanobacteria* like *Arthrospira* and three members of the *Chloroplast* order, as well as genera with various metabolism types such as *Methylibium, Hydrogenophaga* and *Desulfomicrobium.* In addition, it is worth mentioning that from the 3,801 bacterial ASVs detected, 6.5% and 58.3% could not be classified within any known phylum nor genus respectively. Particularly, *Halomonas* and *Marinobacter* are culturable bacteria and seem to be recurrent in the SH, as we reported before (29). Nonetheless, H4 community is by far the most diverse one, according to the Observed, Shannon, Chao and Simpson diversity indices, which were quite heterogeneous between samples (Supplementary Figure S2). Specifically, the phylogenetic diversity showed that the H0 community has a higher number of distant taxa regarding the other communities under study.

As a whole, the beta diversity analysis shows that the dispersion patterns among the communities correlates with what was observed in terms of composition. Hence, the taxonomic composition of H3, H4 and H5 metagenomes are more similar between each other and distinctive from H0 and H1, producing three very well-defined groups (Figure 2B). This is contrasting to the obtained alpha diversity results, suggesting that their members could differ in abundance. Thus, confirming the great level of structure among the communities, considering the total bacterial abundance and taxa diversity. Moreover, the distribution of most abundant taxa among the five communities could explain this segregation. Furthermore, we investigated to what extent the microbial community structure was explained by the environmental factors and found that many variables exert a significant influence over the structure and distribution/grouping, among which arsenic appears to be a main driving force shaping these communities. Besides, salinity along with arsenic separate the H3, H4 and H5 communities, while H0 and H1 segregation is driven by altitude and pH respectively. Nonetheless, for the temperature we have to consider that the observed variation may be the result of the normal daily cycle with respect to the time when samples were taken.

The co-occurrence analysis allowed us to infer possible interactions among complex microbiomes. The network was composed by 112 ASVs having at least one significant correlation, graphically represented as a link between taxa. Total correlations were 155, being 115 positive and 40 negatives. Furthermore, the co-occurrence analyses showed three distinct separated networks within these microbial assembly. However, there is a big main group that comprises most taxa. As expected, the dominant phyla, *Proteobacteria* and *Bacteroidetes*, are among the nodes with a highest degree of interactions (Figure 2C). It is worth mentioning that most of the relevant (highly connected) taxa identified in the networks are little described or studied. Overall, there is the prevalence of direct or indirect relationships among taxa, and this is suggestive of complementing functions in order to maintain the ecosystem. Also, there are few negative correlations that would suggest little niche overlap or competition among taxa.

### Functional approach of SH communities reveals metabolic specialization

The five SH metagenomes we analyzed represent the surface sediment of 5 different sites with a great variation in arsenic content (from 9 to 321 mg/Kg). The 337.5 million quality-controlled reads were co-assembled and yielded a total of 994,545 contigs (≥ 1,000 bp) with 2.38 millions genes, which were used to generate abundance/distributions and functional profiles of the 5 studied communities (Figure 3; Supplementary Table S2). Of those contigs, 148,394 (≥ 2,500 bp) were hierarchically clustered to then be profiled by read-recruitment of the data from the five communities (Figure 3A). The resulting patters are highly variable, considering the contigs detection in each metagenome. In particular, metagenomes from H1 and H5 are the most contrasting ones, as they respectively have 18.78% and 78.40% presence of all available contigs in the SH. Moreover, the rest of the samples also have diverse percentages of representation (H0: 25.00%; H3: 72.08% and H4: 48.33%), and their clustering by the read recruitment profiles correlates with the taxonomic profiles presented previously; being H3, H4 and H5 more similar to each other, and H0 and H1 more distanced from each other and to the others as well. Contrary, the functional profiles seem to have a higher level of convergence between samples in a broad way (Figure 3B). Furthermore, the more enriched categories (SEED Subsystems 1) are associated with metabolism (amino acid derivatives, carbohydrates, protein metabolism, DNA metabolism, cofactor, vitamins, prosthetic groups and pigments) which could be evidence of the needed versatility and adaptability for bacteria to thrive in these harsh environments. Also, the stress response, membrane transporter, cell wall and capsule categories, which are related to the ability to thriving capabilities as well.

**Figure 3.**
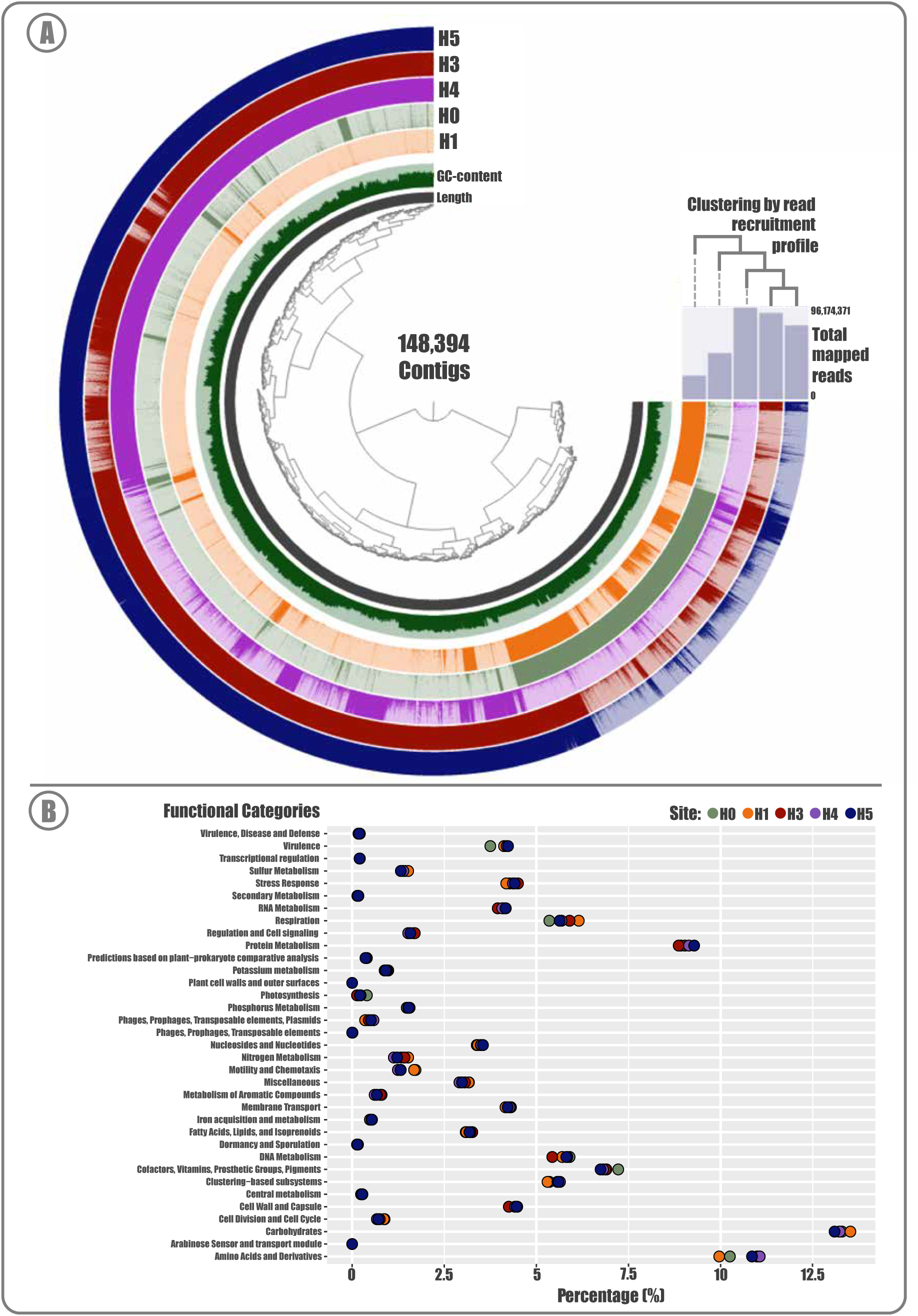
Salar de Huasco metagenomes. **A)** Circular map representing the universe of contigs detected in the five SH sites: the central tree represents the contigs organization on Ward’s linkage with Euclidean distances, the seven circle layers (from bottom-up) represent, for the corresponding contig: its length, GC-content, and presence on the five metagenomes. The top-right bars represent the number of mapped reads for the corresponding metagenome and the dendrogram their clustering by read-recruitment profile. **B)** Patterns of functional potential for each metagenome, according to the presence and abundance of the SEED database of metabolic pathways and functions: subsystems at level 1. The circles represent the percentage value for the corresponding category in each metagenome (defined by colors).

Nonetheless, statistical testing showed that the differences presented by the metagenomes in most categories were significant (Supplementary Table S3). Particularly, the five SH communities showed particular differences in some categories such as arsenic resistance (SEED Subsystems 3), which follows the same tendency previously mentioned; where H0/H1 metagenomes are more similar regarding H3/H4/H5, with 0.21 - 0.22% and 0.26 - 0.27% of reads recruited by this category, respectively. Other significant categories among the communities were carbon, nitrogen, and phosphate metabolism; Zinc, Nickel, Cobalt, Iron and Manganese transport; osmotic stress; circadian clock in *Cyanobacteria* and the Calvin Benson cycle. Whereas, when assessing the ability of these bacterial communities to carry out necessary reactions to sustain some biogeochemical cycles, we found that S, N and CH_4_ showed significant scores (MEBS analysis) for all five metagenomes; implying that the necessary metabolic pathways and machinery is present and with proper completeness (Supplementary Figure S3). Nevertheless, a wide difference is present in the CH_4_ cycle, which is much more enriched in the H1 and H0 communities; especially in the Methane oxidation, Methanol, Methanogenesis and *mcrABC* (markers) pathways. Also, a small decrease in H1 sulfur is also observed.

### Arsenic expulsion is the main mechanism to thrive in the SH

To gain a broader view on the communities’ functional potential related to arsenic metabolism or resistance/tolerance we determined the abundance of the genes belonging to the known mechanisms (Figure 4). Again, the abundance pattern of these genes correlates with the observed tendency, being more related those H3, H4 and H5 sites, distant from H0 and H1. Moreover, although marker genes of arsenic methylation, reduction, oxidation, respiration, and expulsion mechanisms are present in all sites, most are significantly more abundant in the H3, H4 and H5 sites. This may be due to the much higher arsenic concentration in the sediments of these sites. The *arsR* regulator, the *arsC* reductase and the *acr3* pump were the most abundant ones, which would indicate that the As(V) reduction and subsequent As(III) expulsion would be the most common strategy used by the SH inhabitant bacteria. Moreover, *arsM* was also among que most abundant genes, particularly enriched in the H3, H4 and H5 metagenomes. Whereas, genes related to oxidation and respiration mechanisms are less abundant and present an interesting distribution; being those related to oxidation (*aoxA, aoxB, aioB*) are enriched in H3, H4 and H5, contrary to those related to respiration (*arrA, arxR*) are enriched in H0 and H1.

**Figure 4.**
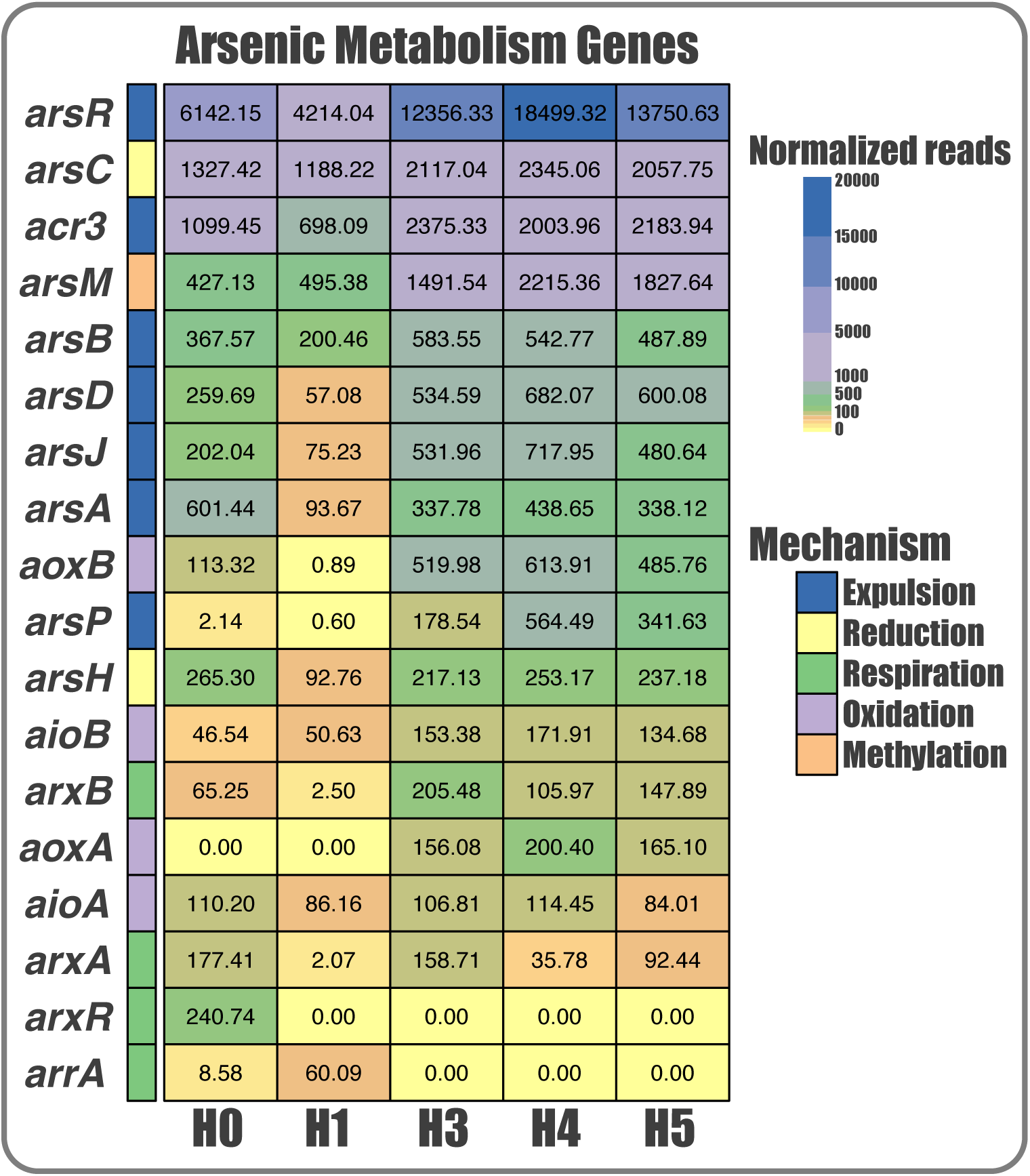
Distribution and abundance of As metabolism genes in the Salar de Huasco. Heatmap shows the (normalized) number of reads that aligned against the corresponding protein identified in each sample, according to the color scale. Genes are grouped by colors, representing the 5 As response mechanisms.

Overall, the proportional distribution pattern of the mechanisms in the 5 metagenomes had some similarity and is quite constant (Figure 5). Particularly, the expulsion mechanism as a whole was the most abundant one for all the sites covering around 75% of all the sequences (Figure 5A); observing the greatest differences in oxidation and respiration mechanisms, as we stated before. Additionally, respiration genes are more present in H0 and reduction in H1. Interestingly, the pattern between the five metagenomes seems to be independent of the amount of detected sequences and the arsenic concentration of the site. Furthermore, comparing the abundance of particular arsenic efflux pumps (from the expulsion mechanism) we can observe that, indeed, most of the sequences correspond to *acr3* (Figure 5B); followed by *arsB* and *arsJ*. Notably, *arsP* which is an efflux permease that confers resistance to organic arsenics (roxarsone and methylarsenite) was more abundant in the sites with higher arsenic concentration. Moreover, the higher *acr3* variants and abundance could be due this pump is present in a wider number of bacteria phyla, covering most of the found diversity (Figure 5C). Also, this efflux pump seems to be more ancestral, regarding *arsP* and *arsJ* which are only present in *Proteobacteria* and in more recent branches.

**Figure 5.**
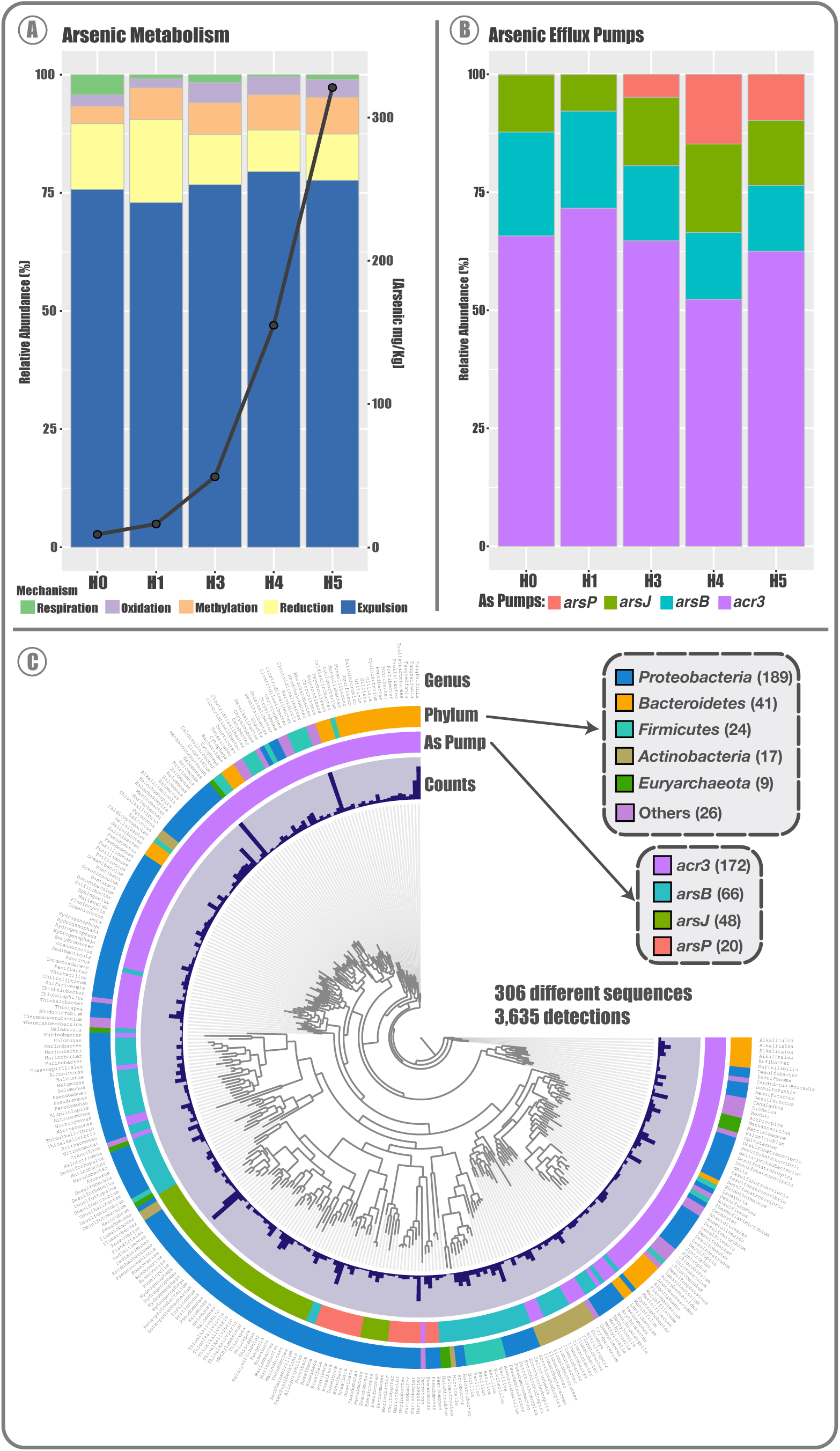
Arsenic metabolism in the Salar de Huasco. **A)** Distribution and abundance of detected genes related to arsenic (As) metabolism in the five SH metagenomes; Stacked bars represent the proportion of all the genes grouped by mechanism type based on relative abundance (%), and the line represents total As concentration in each corresponding site (mg/Kg of sediment). **B)** Stacked bars represent the proportion of all detected As efflux pumps in the five metagenomes, based on genes relative abundance (%). **C)** Phylogenetic analysis of 306 non-redundant sequences of arsenic efflux pumps detected in the SH, layers surrounding the phylogenomic tree indicate: detection level, type of efflux pump, taxonomy, the phylum level, and the species.

### Novel genomes from SH belong undescribed genera

The binning the process reconstructed 195 bins (which were manually curated and evaluated for completion and redundancy) to gain insights of non-culturable bacteria. This resulted in 19 metagenome-assembled genomes (MAGs) that met the completion ≥ 80% and redundancy ≤ 10% criteria (Figure 6); that clustered 4.99% of the contigs in the metagenome profile database. The MAGs taxonomic affiliation resulted in 1/19 belonging to the Eukaryota domain and the 18/19 remaining to the Bacteria domain, distributed in 4 different phyla (9 *Proteobacteria*, 7 *Bacteroidota*, 1 *Cyanobacteria* and 1 *Gemmatimonadetes* (Figure 6A). We only were able to assign 22% (4/18) of the Bacteria MAGs to previously described genera, suggesting a great diversity of novel species that are yet to be described. Also, MAGs contigs number was very variable from 208 to 626 among the Bacteria (Eukaryota being substantially larger: 4,174), the same tendency was observed for the GC-content (36.1% - 69.4%). On the other hand, most of the reconstructed MAGS were represented or detected in the H3, H4 and H5 metagenomes, showing differential abundance patterns in each one (Figure 6B; Supplementary Table S4).

**Figure 6.**
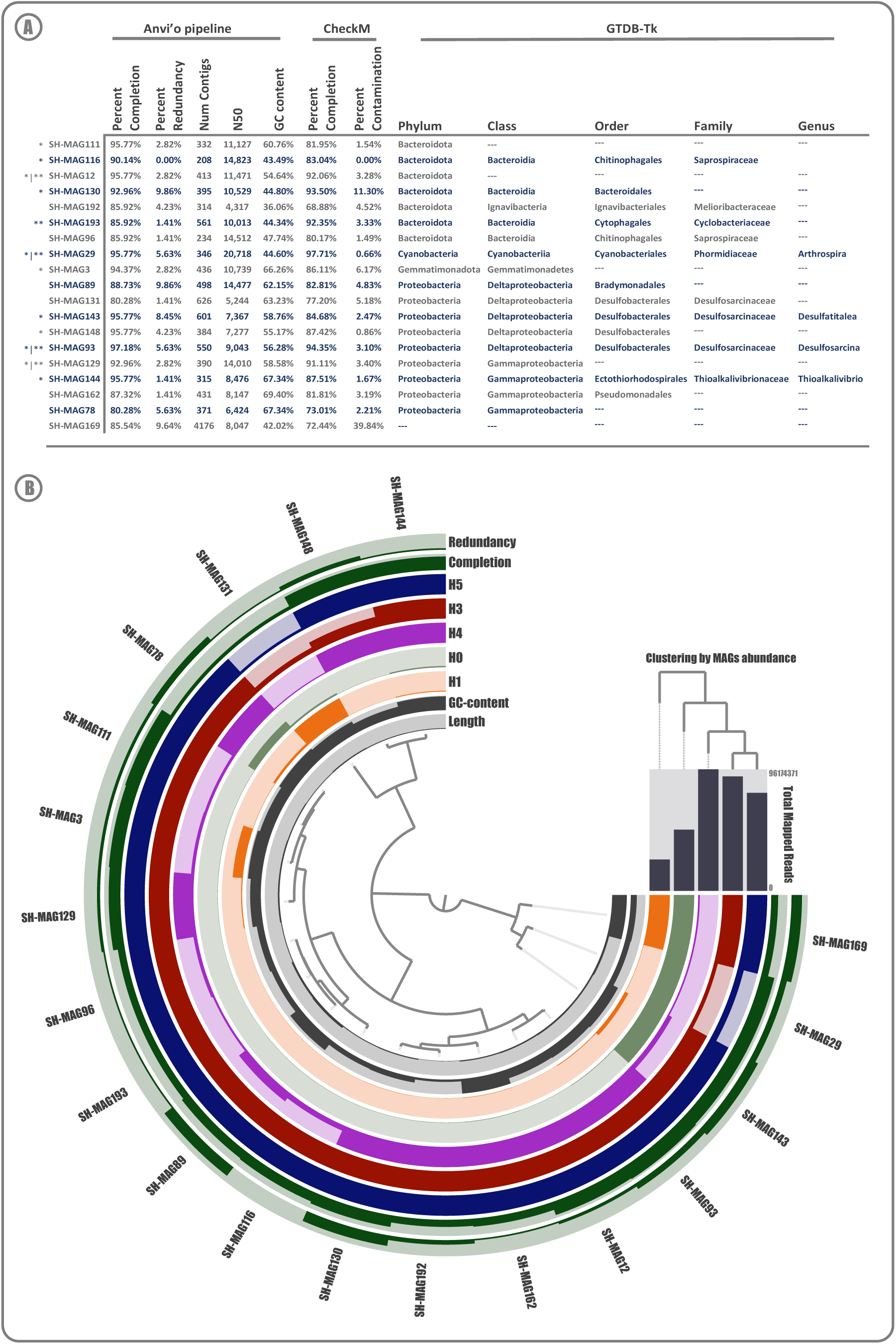
Salar de Huasco Metagenome-Assembled Genomes. **A)** Genomic features summary of the SH 19 MAGs, quality indices, and taxonomic affiliation (the results of *Anvi’o pipeline and/or **CheckM meet the quality standard of completion ≥ 90% and redundancy/contamination ≤ 10% (according to 125). **B)** Presence and abundance of the 19 MAGs on the five SH metagenomes: the central tree represents the MAGs organization on Ward’s linkage with Euclidean distances, the seven circle layers (from bottom-up) represent, for the corresponding MAG: its length, GC-content, and abundance on the five metagenomes. The top-right bars represent the total number of mapped reads for the corresponding metagenome and the dendrogram their clustering by absence/presence profile.

### Novel genomes from SH are mostly arsenic reducers

The functional potential of reconstructed MAGs was evaluated globally and particularly in relation to their repertoire of arsenic associated genes (Figure 7). For these evaluations, only the 12 MAGs that meet the quality standard of completion ≥90% and redundancy/contamination ≤10% were considered. Therefore, a total of 267 KEGG modules were detected among the MAGs, presenting variable absence/presence and completion patterns (Figure 7A; Supplementary Table S5). This variation reflects the MAGs taxonomic affiliations, being all the *Bacteroidota* (MAG111, MAG12, MAG116, MAG130 and MAG193) clustered together, as well as the *Deltaproteobacteria* (MAG148, MAG143 and MAG93) and *Gammaproteobacteria* (MAG129 and MAG144). On the other hand, the only *Cyanobacteria* (MAG29) is close to the *Gammaproteobacteria* and the only *Gemmatimonadota* (MAG3) is close to the *Bacteroidota.* Particularly, the MAGs presented a considerable number of genes regarding arsenic metabolism, where the same tendency shown before was observed; with a dominant presence of genes related to As expulsion (*acr3, arsA, arsJ* and *arsR*) throughout all taxa, followed by reduction (*arsC* and *arsH*) and methylation (*arsM*) (Figure 7B, Supplementary Table S6). Interestingly, *arsJ* was only detected in one *Proteobacteria*, which agrees with its lower abundance found at the metagenomic level in the SH. While, *arsH* was only detected in the *Cyanobacteria,* which could also reflect the abundance of this taxa and this gene globally. In addition, the absence/presence patterns among the MAGs seems to be group-specific, broadly associated by phyla.

**Figure 7.**
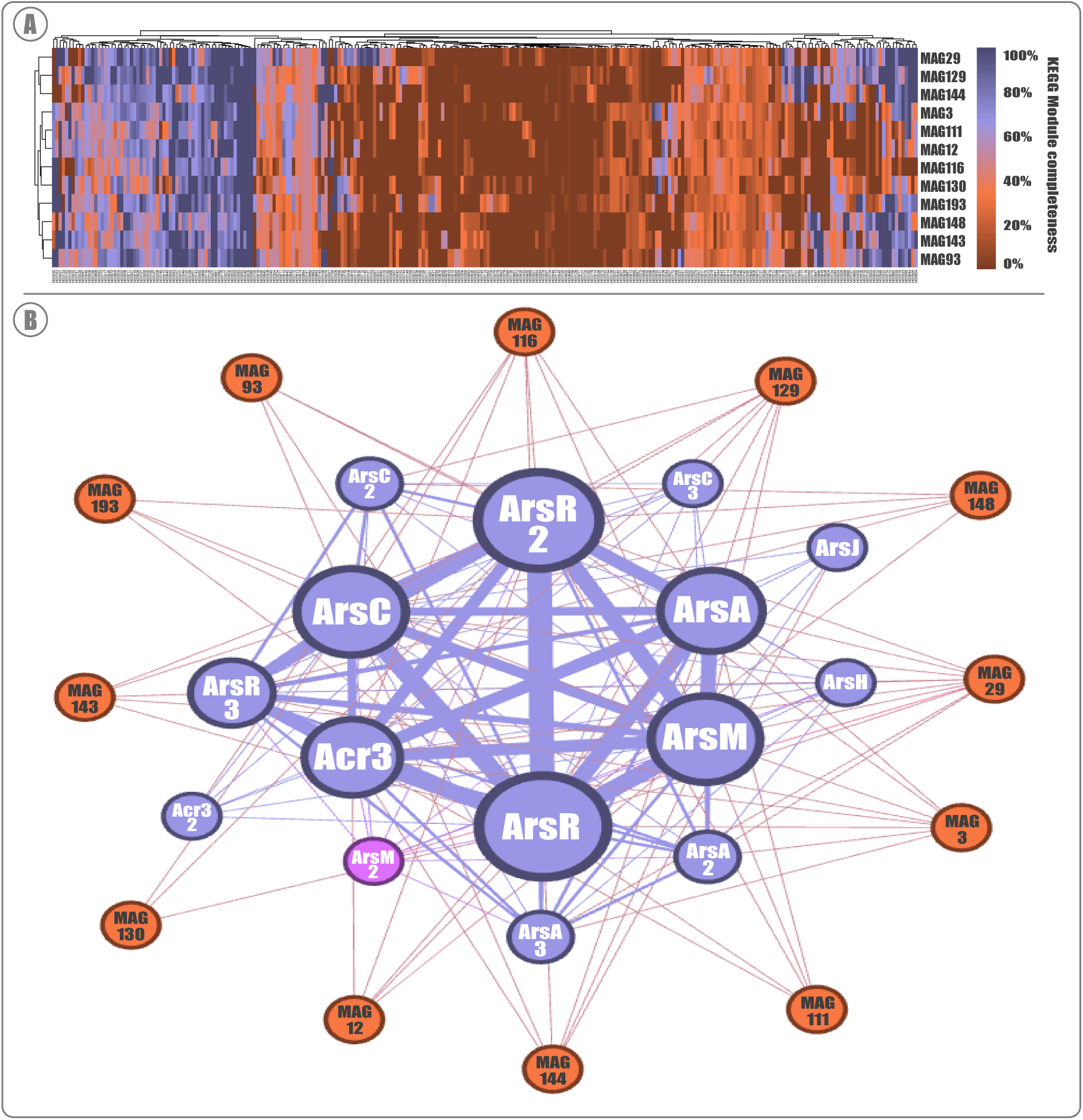
MAGs functional potential profiles: global and arsenic specific. **A)** Metabolic capabilities of the 12 selected MAGs according to KOfam (KEGG Orthologs database); the heatmap shows the KEGG Modules completition displayed by the color gradient for given module in each corresponding MAG. **B)** Functional network analysis of the 12 MAGs according to the presence and abundance of detected arsenic metabolism genes; size of gene nodes represents the level of detection, and the size of the edges represents the correlation level. Blue nodes represent genes, and orange nodes represent MAGs.

## DISCUSSION

The Salar de Huasco has an important spatial variation of arsenic in sediments, which we have described as a gradient from north to south (Figure 1) (9); being consistent with the widely reported heterogeneity for this area and the impact of extreme conditions over the inhabitant organisms (30). Previous studies have detected variations in conductivity, organic matter, and dissolved oxygen even in geographically closed areas (6, 7, 31, 32, 33). In aquatic ecosystems, the quantity of trace elements and physical-chemical states of some elements, are related to their circulation/interaction, and with the mechanisms of corresponding living organism (34). Salar flats as SH have the peculiarity of containing more than 50% of CaCO_3_ and high levels of arsenic, which is very available and highly mobile. This is because a fraction of the arsenic in sediments lies as soluble salts (Na_2_HAsO_4_); it also could be associated to the calcium carbonate (CaHAsO_4_) and, adsorbed into Fe and Al oxides (35). The metabolism of inhabiting microbial communities or the activity of primary producers could be the source of the diversity of abiotic properties (36). It is hypothesized that this spatial and functional heterogeneity is the cornerstone of resilience in these extreme ecosystems. We must consider the presence of high concentrations of arsenic as a selective pressure, which could have a modulating effect on the composition of the H3, H4 and H5 communities (grouping them together and separating them from the rest).

The five communities described from the SH metagenomes are dominated by *Proteobacteria, Bacteroidetes,* and *Cyanobacteria* (Figure 2), similar to previous reports for other Altiplanic environments (Socompa volcanic Lake in Argentina and La Brava Lagoon - Salar de Atacama in Chile) mostly carried out by 16S rRNA amplicon sequencing (37, 38). Specifically, the same pattern has been reported previously for the SH (3, 32). However, the proportions are different, probably because the shotgun metagenomic sequencing used in this study is more sensitive, which is evidenced by the alpha diversity indiceś magnitude. Nevertheless, these values are also higher than those reported by other metagenomic studies from similar areas (39).

Moreover, there is evidence of *Proteobacteria* enrichment in places with different arsenic concentrations (12), as many microorganisms described in this group are capable of interacting and tolerate arsenic. Some examples are *Acidithiobacillus* and *Desulfovibrio* genera, which can solubilize arsenic from solid compounds or precipitate it by coupling arsenic and sulfur reduction, respectively (10, 40). The phylum *Bacteroidetes* is widespread and commonly found in hyper-saline wetlands and microbial mats (38, 41, 42, 43, 44, 45). Furthermore, *Cyanobacteria* have fundamental roles in any community they belong to, as primary producers and participating in bio-weathering and matrix transformation processes (46). Also, they are usually abundant in places exposed to sunlight (47). Besides, previous reports have proposed the cyanobacterial communities in SH are unique (6). In addition, many recurrent representatives of these communities (*Marinobacter* and *Halomonas*) have been cultured and isolated in laboratory conditions (29). Particularly, we isolated *Exiguobacterium* strains from these communities to describe the mechanisms responsible for their high arsenic resistance (9, 48, 49).

On the other hand, a high percentage of unidentified genera have also been reported in the Chilean Atacama Desert, where those reach 66% putatively novel taxa belonging to that bacterial “dark matter” (50), this is a direct consequence of database shortage, reinforcing the need to keep exploring these unique environments and describe the novel, highly adapted microorganisms that remain unknown to science and that are of fundamental importance to the maintenance of these ecosystems. The co-occurrence networks show clear evidence of overlapping among the bacterial taxonomic composition of the metagenomes or “core microbiome”, mainly composed of unknown and little-studied/described taxa. As an interaction example, we have *Seohaeicola* and *Anoxynatronum*, belonging to an isolated in the network involved in nitrogen cycling as autotrophic and heterotrophic taxa. From the co-occurrence networks, we can infer that this is a very fragile and susceptible ecosystem as there are no keystone nodes detected (51).

At the lower available taxonomic rank, we found some bacteria that are typical for extreme environments, such as *Thiobacillus*, a well-known sulfur-oxidizing bacterium, and *Desulfotignum* an anaerobic group known to be sulphate-reductors, as well as other less studied groups. The most abundant belong to the *Roseovarius* genus, an aerobic, non-photoautotroph bacterium; *Rhodohalobacter* is a facultative anaerobic, moderately halophilic group, and *Truepera* comprises aerobic quimio-organotropic alkaliphile bacterium. This is evidence of the lifestyle diversity that these microorganisms have, suggesting a cooperative and specialized community. Likewise, this is in coherence with our other findings, the enrichment of photosynthetic organisms in sites H0 and H1, as well as *Proteobacteria* as the most abundant group in all the other sampled sites. Other similarities are evidenced, as the genus *Methylibium* is only found on the H0 site, where the methane-related metabolism function was found to be enriched in the MEBS analyses. In addition, the importance of some of these abundant taxa is highlighted, as they are structural parts in the co-occurrence network, namely *Desulfomicrobium, Thiobacillus,* and *Brumimicrobium*.

The functional potential of the five SH communities showed similar patterns in general, with particular differences in some categories following the same association pattern described before (Figure 3). Comparing each site, we found significant differences in the enrichment of some important functional categories related to the maintenance of critical geochemical cycles, including carbon, nitrogen, and phosphate metabolism, stress resistance, transport of Zinc, Nickel, Cobalt, Iron, Manganese and osmotic stress. Notably, the two functions with statistical differences between almost all communities were the Circadian Clock in *Cyanobacteria* and the Calvin Benson cycle, suggesting differences in the primary production and CO_2_ fixation. The Circadian Rhythm is directly influenced by several abiotic conditions, such as temperature and atmospheric pressure (52). Moreover, the Calvin Benson cycle was the only metabolic pathway involved in CO_2_ fixation reported in an endolithic halite metagenome analysis, underlining its importance for microbial communities living in these extreme conditions (53). This agrees with the MEBS analysis results, where the carbon cycle relevant reactions presented a greater variation between the metagenomes. Particularly, carbon flux control in these communities seems to be different especially in regard to methanogenesis processes, as the far greater detection of specific markers in H0 and H1 could account for a high presence of methanogenic species in these two communities in comparison to the other metagenomes (54). This could be related to the S decrease in H1, implying that methanogens outcompete sulfate reducers in this community (55).

Furthermore, among these significant differences, arsenic resistance is enriched in the metagenomes from the sites with the highest As concentration, which was expected and has been reported previously (56, 57). Particularly, the distribution and abundance of arsenic-related genes in the five SH communities showed significant differences among the communities (Figure 4), as the presence of arsenic affects the dynamics of inhabiting microorganisms (18, 58). Therefore, arsenic would promote the selection and abundance of tolerant microorganisms (59, 60). Our results agree with this premise since the abundance of all these genes increases following the gradient of arsenic concentration among the five sites, indicating a different dynamic in each niche, even though there are geographically closely located.

Overall, we found that most of the detected genes related to arsenic belong to expulsion and reduction mechanisms (*arsR, arsC,* and *acr3*), which correlates with what was reported in the Argentinian Altiplano (22). The transcriptional repressor ArsR was the one with the highest abundance detected; as this protein participates in the regulation of different arsenic pumps, namely ArsB, ACR3, and ArsP (61, 62); and is also reported to be in operon conformations along with ArsC or ArsM (63). Hence, it was expected to be the most abundant and broadly distributed. Moreover, to better assess the proportion of the arsenic mechanisms found in SH, we separated the genes related to As expulsion out of the cell from those related to reduction mechanism (usually reported together). This is due to the fact that gene clusters including *arsB, arsP, arsK*, *arsJ,* and *acr3* are commonly found/reported independent of arsenate reductase *arsC* (9, 62, 64, 65). Also, arsenic methyltransferase (ArsM) was the next most abundant protein, and particularly enriched in H3, H4, and H5, suggesting this protein is part of a complementary mechanism used to thrive in sites with high arsenic concentrations (66).

Even though proteins related to oxidation and respiration of As are less abundant; their distribution is interesting, which could be due to niche-specific conditions. Despite these mechanisms are somewhat known and recurrent, much remains unclear, mostly regarding their function and interconnection with the bacterial central metabolism (67, 68). This problem is exacerbated when working in unexplored environments, where unknown or unclassified organisms abound. Besides, the amount of misclassified and/or unclassified protein sequences in databases is a major blockage for these investigations. Nonetheless, this is fully acknowledged and is being addressed; thus, helping to generate important findings (69, 70, 71, 72). For example, our group was able to manually identify the ArsP and ArsK transporters, as well as a possible arsenic respiration system, additionally a missing arsenic methylase was evidenced in *Exiguobacterium* genomes using a combination of bioinformatic tools (9). Nevertheless, database shortage and the classification by homology problems must always be considered when gene- or metabolism-detection is being carried out.

Broadly, the most abundant mechanism found in the SH was arsenic expulsion, with around 75% of all the arsenic-related sequences (Figure 5). Nonetheless, by merging the expulsion and reduction mechanisms, we would get results consistent with previous reports for paddy, contaminated, and natural arsenic-rich soils (around 85%) for the reduction mechanism (56). Likewise, here we report that mechanism proportions are relatively equivalent between the five SH sites, and those are independent of the total sequence abundance as well as the arsenic concentration. Whereas respiration and oxidation mechanisms showed some variation, which could be reflecting the taxonomic composition of the communities. Furthermore, we also found that the Organoarsenical Permease *arsP* was detected in the higher arsenic concentration sites. Therefore, this may be a counteraction against a possible greater production of these highly toxic organic species at these sites, where consistently the Arsenite Methyltransferase (*arsM*) gene was enriched as well. Thus, these markers could be proposed as an arsenic contamination bioindicator like *acr3* (73). Additionally, the distribution and phylogenetic relationships of arsenic pumps among the metagenomes revealed distinct groups, which is consistent with those previously reported: I) ArsB - Acr3 [ion/BART], II) ArsP [Permease]; III) ArsK - ArsJ [MFS] (65). It has been hypothesized that the divergence presented by the arsenic resistance mechanisms originated from Earth’s geological changes to adapt for a particular function or As emergent species (74). Besides, we found that *Proteobacteria* possess a broader gene repertoire for arsenic response, which has also been described before (22).

The twelve reconstructed high-quality MAGs provide information about the abundance and distribution of undescribed or unknown microorganisms (Figure 6), as well as genomic insights into widespread arsenic metabolism. Most of these are affiliated within *Proteobacteria* and *Bacteroidetes* phyla, reflecting what we described at the community level. As stated before, the recovery of MAGs complements decades of cultivation and PCR-survey efforts by providing information about taxa missing in culture collections (Candidate Phyla Radiation), improving our understanding of microbial communities (75); particularly those inhabiting extreme environments. Nonetheless, four MAGs were classified at the genus rank: *Arthrospira*, a *Cyanobacteria* with great commercial interest as a pigment source and to make spirulina supplements; *Thioalkalivibrio:* an aerobic sulfur-oxidizer; *Desulfatitalea* and *Desulfosarcina*: two anaerobic sulfate reducers (76, 77, 78).

In the same way that these MAGs only cover the most abundant phyla of SH, we only detected genes that belong to the most prevalent arsenic resistance mechanisms on them (Figure 7). Thus, supporting expulsion, reduction, and methylation as the most recurrent widespread mechanisms in the SH to cope with the high arsenic concentrations. Conversely, no arsenic oxidation and/or respiration-related gene was detected on the recovered MAGs. This could indicate that the different ways used to expel the toxic out of the cell would be more straightforward and direct as a first line of defense against arsenic; while the oxidation and respiration would involve a coupling with the bacteria central metabolism and a larger machinery to benefit from this compound, which seems to be associated with less abundant and highly adapted or specialized bacteria. Also, arsenic oxidation and respiration have been reported in association with nitrate and sulfate reduction, respectively (10, 79). Nonetheless, we need to take into consideration the MAGs completion levels as a possible source for missing genes.

Overall, our results reveal that populations of *Proteobacteria, Bacteroidetes* and *Cyanobacteria* are abundant across wide ecological niches in the SH spanning a challenging ensemble of environmental conditions and physico-chemical parameters, among which arsenic is highly relevant. As the arsenic cycle and the bacterial contribution to it, has yet to be completely understood, more environmental studies are needed, while metagenomic approaches have been shedding light on the possible role of unknown or undescribed microorganisms additional transcriptomic, metabolomic and cultivation approaches will be essential to define these phenomena and its relevance in a global context.

## CONCLUSION

The Altiplano array of ecological niches is a reservoir for microbial diversity, showing great richness in adapted organisms capable of facing these challenging conditions. Particularly, Salar de Huasco is a highly diverse ecosystem, where salinity and As concentration contribute to shaping the community composition, mainly represented by *Proteobacteria* and *Bacteroidetes*. Also, the interaction networks within these communities showed three distinct groups of related taxa, but no “key stone” nodes were found. Nonetheless, little niche overlap was determined. Altogether, these indicate that SH studied niches harbor highly diverse communities, being H1 and H5 the most contrasting ones. Moreover, the most abundant arsenic-related genes found in these communities indicate that the As(V) reduction and subsequent As(III) expulsion would be the most common strategy used to detoxify the cell of arsenic; furthermore, regarding to the expulsion pumps, the most abundant were Acr3, followed by ArsB; however, in sites with high As concentration ArsP begins to be enriched. In addition, 12 high-quality, non-redundant MAGs were reconstructed from the metagenomes; those represented the dominant diversity detected across the communities as well as the metabolic variability and the presence of marker-genes related to the most recurrent As resistance mechanisms (expulsion, reduction and methylation). Finally, in order to further elucidate the strategies and relationships between the microbial taxa and among biotic and abiotic components within the ecosystem, further multidisciplinary studies are required, as well as the use of the ever evolving NGS approaches with the increasing database information to better understand the evolutionary process of adaptation to the extreme conditions presented by these unique ecosystems.

## MATERIALS AND METHODS

### Study Area and Sampling

Salar de Huasco National Park (SH) is an area located on the Chilean Altiplano that is known for its spatial heterogeneity, as well as great biodiversity and physicochemical characteristics. This ecosystem is mostly composed of streams, salt crusts, peatlands, and shallow (permanent and non-permanent) lakes with salinity and arsenic gradients from north to south (9). We sampled surface sediments (to a depth of 5 cm) in sterile tubes from five different sites (H0 to H5 previously described in (6)) by duplicate during fieldwork done in June 2018; those samples were kept and transported in a cooler until stored at −20°C for subsequent DNA extraction. Physicochemical parameters like temperature, salinity, and pH were recorded (HI 98192 and HI 2211 - HANNA Instruments) *in situ* (Figure 1).

### DNA extraction and high throughput shotgun sequencing

Total DNA was extracted from the sediment samples from each SH site using the DNeasy PowerSoil Kit (Qiagen Inc., Hilden, Germany) following manufacturer’s instructions. DNA integrity, quality, and quantity were verified through 1% agarose gel electrophoresis, OD_260/280_ ratio, and fluorescence using a Qubit® 3.0 Fluorometer along with the Qubit dsDNA HS Assay Kit (Thermo Fisher Scientific, MA, USA). Next, Paired-end (150bp) libraries were constructed for each sample in duplicates, at the Centro de Biotecnología Vegetal, Universidad Andrés Bello (Santiago, Chile) using the TruSeq Nano DNA Kit (Illumina Inc., CA, USA.) following the TruSeq Nano DNA Sample Preparation Guide 15041110 Rev. D. Libraries were sent for sequencing at Macrogen Inc. (Seoul, Korea) on a HiSeq 4000 platform (Illumina Inc., CA, USA). Then, raw data was evaluated using FastQC v0.11.8 (80) for quality control, adapters were removed from the reads of all samples using Trimmomatic v0.30 (81) and the filtering and trimming (length ≤ 100bp, Ns = 0, and Q ≤ 30 thresholds) was performed with PRINSEQ v0.20.4 (82). The whole raw data sets have been deposited at DDBJ/ENA/GenBank under the Bioproject: PRJNA573913.

### Taxonomic profiling analysis

Quality-controlled reads for each sample were profiled using the phyloFlash pipeline (83) to obtain all reads that align with the bacterial SSU rRNA (small-subunit rRNA) SILVA v138 database (84). Later, these sequences (FASTQ files) were processed using R v3.5.2 and RStudio v1.1.463 (85, 86) following the DADA2 v1.16.0 R package pipeline (87) to infer amplicon sequence variants (ASVs) present in each sample. Briefly, after dereplication, denoising, and paired reads merge steps, the ASV table was built with 97% clustering, the chimeras removed, and the taxonomic was assigned against the Silva v138 database (84) using DADA2 Ribosomal Database Project’s (RDP) naive Bayesian classifier (88). Then, data was normalized by variance stabilizing transformation using the R package DESeq2 v1.28.1 (89). Also, a multiple sequence alignment was created using the R package DECIPHER v2.16.1 (90) to infer a phylogeny with FastTree v2.1.10 (91). Furthermore, a phyloseq-object (containing the ASVs, taxonomy assignation, phylogenetic tree, and samples meta-data) was created using the R package Phyloseq v1.32.0 (92); in order to calculate the alpha diversity indexes, along with btools v0.0.1 R package. Also, beta diversity (PCoA - Bray Curtis distance with environmental variables fit) was calculated using the R package ampvis2 v2.4.5 (93), and visualizations were generated with ggplot2 v2.2.1 (94) R package.

### Co-occurrence networks

We used the same phyloseq object, which was agglomerated by best hit using the microbiomeutilities v1.00.11 R package (95); filtered by tax abundance (0.5% in at least one sample) using the Genefilter v 1.72.0 (96) and Phyloseq v1.32.0 (92) R packages. Then, the co-occurrence network was estimated using the SPIEC-EASI v0.1.4 R package (SParse InversE Covariance Estimation for Ecological Association Inference) (97), using neighborhood selection model (lambda.min.ratio=1e-2, nlambda=20, and 50 replicates parameters). Finally, the network was visualized using the ggnet2 function of GGally v1.5.0 R package (a ggplot2 extension) (98).

### Functional profiling analysis

The patterns of functional potential as subsystems with different specificity levels for each community were determined by the presence/absence and relative abundance of the quality-controlled reads that aligned against the metabolic pathways and functions in the SEED database (99) with SUPER-FOCUS v0.35 software (100), which uses DIAMOND v2.0.6 (101) for fast and efficient alignment. Statistical analysis was carried out in STAMP v2.3.1 (Software package for Analyzing Taxonomic or Metabolic Profiles) (102) using Welch’s t-test to compare all samples.

### Metagenome co-assembly and read mapping

As we need for subsequent analyses: i) a co-assembly (the reads from all samples assembled together: .fasta file), ii) its annotations (.gff, .ffn and .faa files) and iii) an alignment per samples (all the reads from each sample mapped against the co-assembly: .bam files); we proceeded to assemble the quality-controlled reads from all samples using MEGAHIT v1.1.3 (103), with the –presets meta-large option and a minimum contig length of 1 kb. The co-assembly was then evaluated with MetaQUAST v5.0.2 (104) and annotated with Prokka v1.11 (105) using the metagenome mode. Furthermore, we mapped the quality-controlled reads from each sample against the co-assembly using Bowtie2 v2.3.4 (106) and stored the recruited reads (sorted and indexed) as BAM files using SAMtools v1.3 (107).

### Metagenome profiling

We followed the anvi’o v7 pipeline (108). First, we created a contig database with the co-assembed contigs, which uses Prodigal v2.6.3 (109), HMMER v3.3.1 (110) and NCBI COGs (111) to identify genes calls and functionally annotate them (anvi-gen-contigs-database, anvi-run-hmms, anvi-run-ncbi-cogs). Secondly, we profiled each sample’s BAM file against the contigs database to estimate the detection and coverage statistics for each contig (anvi-profile). Then, we combined the profiles in a single merged metagenomic profile database, which uses all individual statistics to compute hierarchical clustering of the contigs (anvi-merge). Finally, we visualized the merged profile on the anvi’o interactive interface, which allows easy exploration and curation of the metagenomes (anvi-interactive). On the other hand, we used MEBS (Multigenomic Entropy Based Score) v1.0 package (112) to evaluate and compare the S, N, O, CH_4_, and Fe biogeochemical cycles on each metagenome using the predicted proteins (annotated .faa file from each individual assembly).

### Metagenome target gene search

To estimate the abundance of genes related to arsenic metabolism, the read counts for each predicted gene of each sample were obtained using the corresponding BAM file and the co-assembly annotated GFF file, with HTSeq-Counts v0.13.5 (113). Also, we constructed a database from the GenBank – “Identical Protein Groups” with all available prokaryotic proteins of arsenic metabolism (Acr3, AioA, AioB, AioR, AioS, AioX, AoxA, AoxB, AoxC, AoxD, ArrA, ArrB, ArrC, ArsA, ArsB, ArsC, ArsD, ArsH, ArsJ, ArsK, ArsM, ArsN, ArsO, ArsP, ArsR, ArsT, ArxA, ArxB, ArxR, ArxS and ArxX) and this was targeted with the co-assembly predicted proteins (.faa file) using CRB-BLAST v0.6.6 (E-value ≥1E^−05^, Identity ≥70% and Query Coverage ≥70%) (114). Finally, matching the hits with the gene counts (normalized by the target gene length and the corresponding library size), we calculated the relative abundance of the interest genes for the five metagenomes. Visualizations were made with R packages ggplot2 v2.2.1 (94) and pheatmap v1.0.12 (115). Furthermore, all the detected sequences that corresponded to arsenic efflux pumps were extracted from the .ffn file and aligned using MAFFT v7 (116). As phylogeny inference was calculated with FastTree v2.1.10 (91) and visualized with the anvi’o v7 interactive interface (108).

### Metagenomic binning

The contigs in the metagenome profile were clustered through anvi’o v7 (108) using CONCOCT v1.1.0 (117) binning program, which adds a bins collection the profile (anvi-cluster-contigs). The resulting bins were evaluated for completion/redundancy, and manual curation/refinement was carried out in the interactive interface (anvi-estimate-genome-completeness and anvi-refine). Finally, the refined bins were displayed in the interactive interface and summarized to obtain all the statistics and files for downstream analysis (anvi-interactive and anvi-summarize).

### MAGs evaluation, taxonomy, and functional estimation

We define MAGs (Metagenome Assembles Genomes) as bins with a completion >80% and redundancy <10%. Then, we used CheckM v1.0.13 (118) for a more robust evaluation. Later, to infer MAGs taxonomy, we used GTDB-Tk v0.3.2 (119) along with The Genome Taxonomy Database (120). Moreover, we estimated the MAGs metabolic potential by evaluating their gene content with anvi’o v7 (108); First, functions and metabolic pathways were annotated to the MAGs using HMM hits from KOfam - KEGG Orthologs (KO) database (121, 122) (anvi-run-kegg-kofams). Secondly, relying on these KO annotations, the metabolic pathways were predicted considering those defined by KOs in the KEGG MODULES resource (123), where a KO represents a gene function, and a module represent a group of KOs that together carry out the reactions in a metabolic path (anvi-estimate-metabolism). The MAGs module completition was visualized using pheatmap v1.0.12 R package (115).

### MAGs target gene search

All MAGs were queried against the same previously constructed database (with the arsenic metabolism genes) using CRB-BLAST v0.6.6 (E-value ≥1E^−05^, Identity ≥70% and Query Coverage ≥70%) (114). Furthermore, the resulting hits matrix was compared and supplemented with the detected KOfams related to arsenic, and the final matrix was used to generate a functional network using Gephi v0.9.2 (124) to connect MAGs and detected genes.

## DATA AVAILABILITY

The whole raw data sets have been deposited at DDBJ/ENA/GenBank under the Bioproject: PRJNA573913.

## AUTHOR CONTRIBUTIONS

JC-S, CP-E, EC-N, FR, and CPS conceived and designed the study. JC-S, JF, FM, FR, and CPS performed the field work. JC-S and CP-E processed the samples, performed the experimental procedures and carried out the bioinformatics analyzes. KM, SM and EC-N contributed with critical bioinformatics advice. CPS, FR, and EC-N contributed with reagents, materials, and analysis tools. JC-S and CP-E interpreted the results and wrote the first manuscript draft. All authors read and approved the final manuscript.

## ACKNOWLEDGEMENTS

We would like to thank the illustrator Florence Gutzwiller for the SH landscape painting (https://spideryscrawl-illustration.webnode.com/). Also, we would like to thank Universidad Andres Bello’s high-performance computing cluster, Dylan (http://www.castrolab.org/), for providing data storage, support, and computing power for bioinformatic analyses. Also, We would also like to thank the MerenLab group (https://merenlab.org/) for their reachability in providing help, advice and solutions related to the use of Anvi’o. In addition, wéd like to thank to Dra. Valerie de Anda (https://valdeanda.github.io/) for the guidance and advice to implement the MEBS software.

## SUPPLEMENTARY MATERIAL

**Supplementary Table S1.** Relative abundance of all detected phyla in the five SH communities.

**Supplementary Table S2.** Statistical values of the SH co-assembly, representing the five metagenomes.

**Supplementary Table S3.** Statistical evaluation of enriched functional categories (SEED subsystem 1) in each of the five communities regarding the rest according to Welch’s t-test.

**Supplementary Table S4.** Indexes of relative abundance and detection of each recovered MAG across the five SH metagenomes.

**Supplementary Table S5.** Completion index of all detected KEGG modules each analyzed MAG.

**Supplementary Table S6.** Presence and copy number of arsenic resistance related genes in each analyzed MAG.

---

**Supplementary Figure S1.**
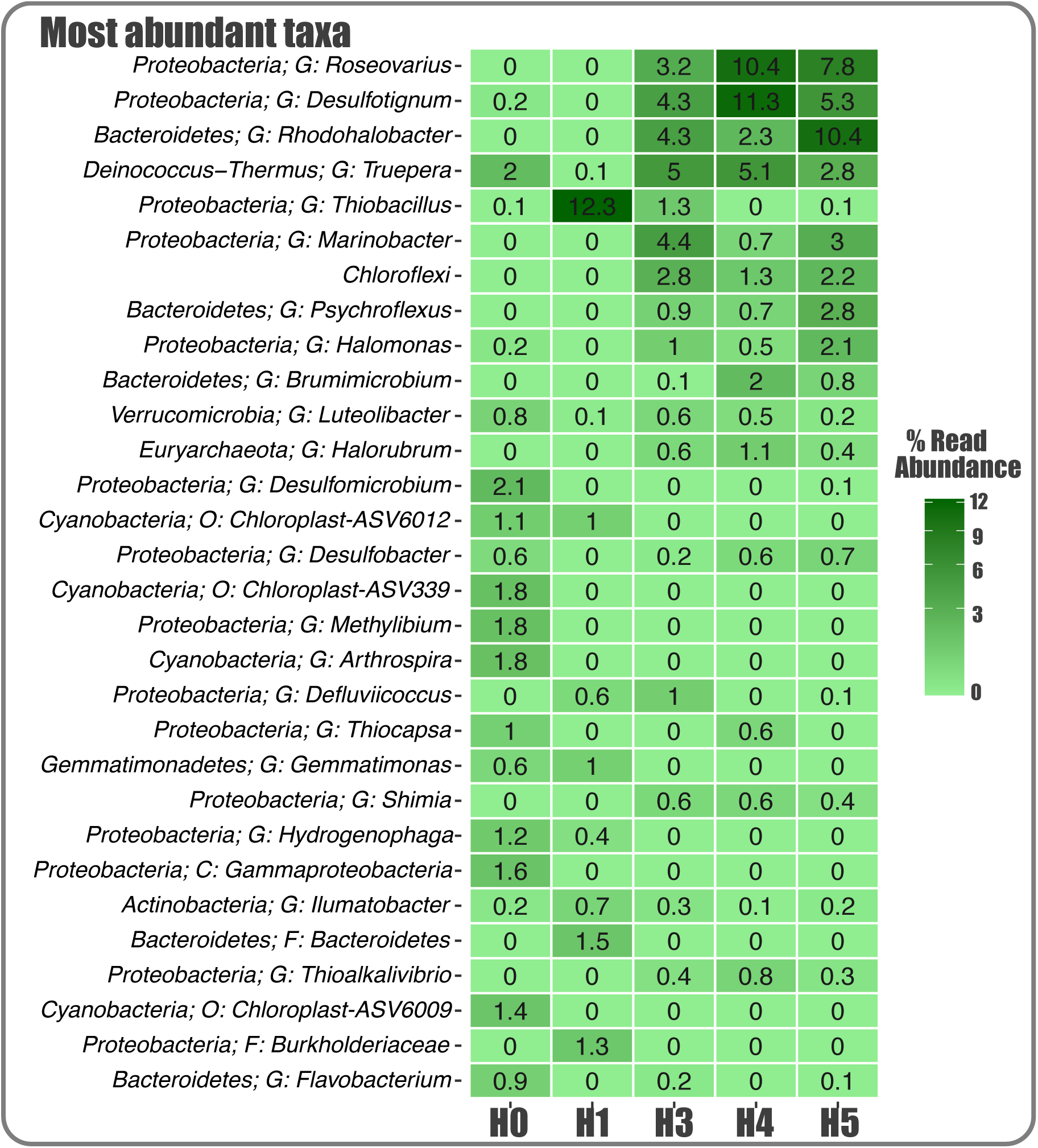
Composition and structure of SH bacterial communities. Heatmap showing the 30 most abundant ASVs (bacterial lineages). Taxonomic is showed at the phylum rank plus the best hit available (C: class; O: order; F: family; G: genus).

**Supplementary Figure S2.**
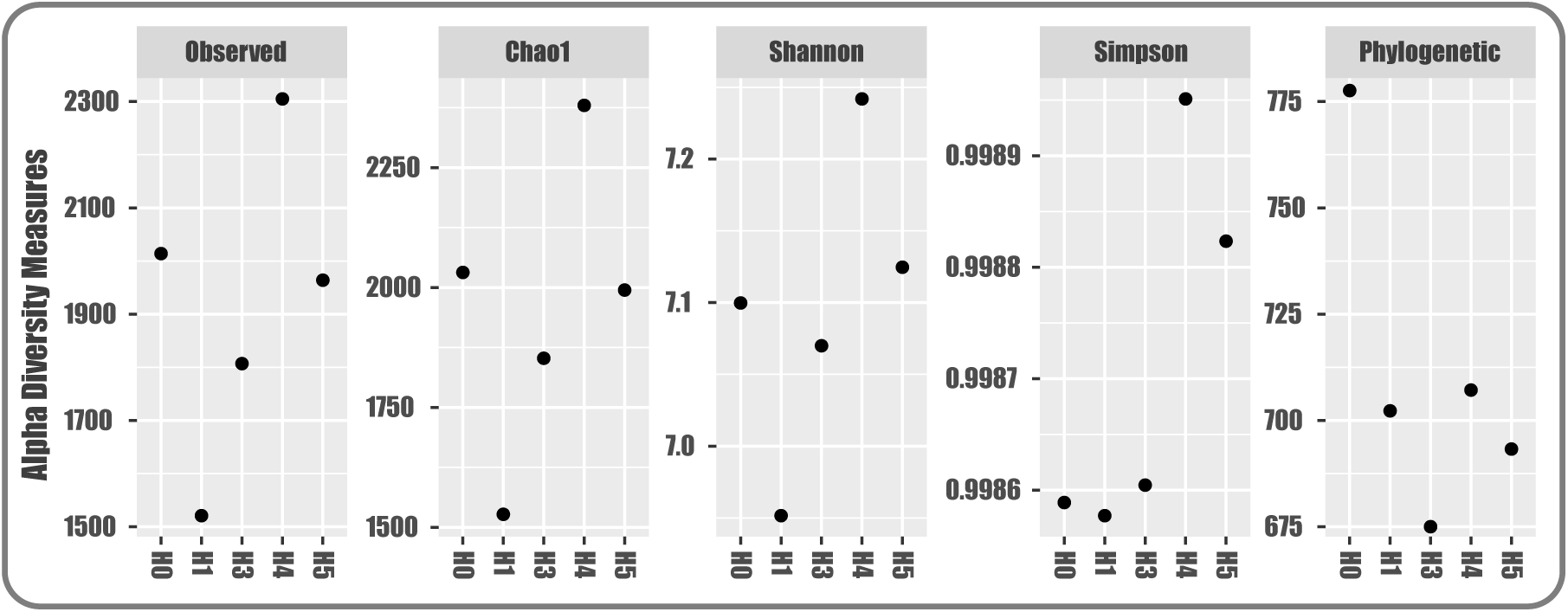
Alpha diversity indices for the five SH communities.

**Supplementary Figure S3.**
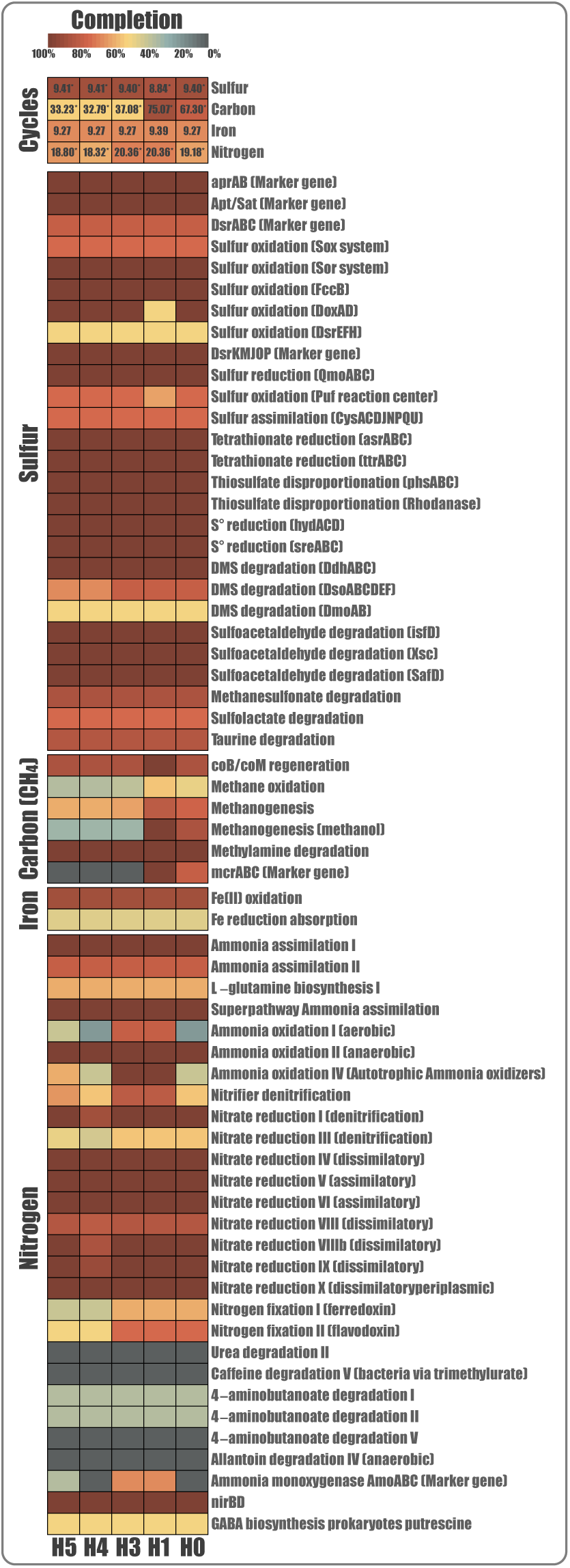
MEBS analysis heatmap displaying the completeness of N, Fe, S and CH_4_ pathways: as a whole (first top section) and particular reactions (bottom four sections). The color gradient shows the percentage of completition for each pathway (from lowest to highest) and the values at the top section represent the corresponding MEBS score (* FDR ≤ 0.01).

## REFERENCES

1. Bahram M, Netherway T, Frioux C, Ferretti P, Coelho LP, Geisen S, Bork P, Hildebrand F. 2020. Metagenomic assessment of the global diversity and distribution of bacteria and fungi. Env Microbiol 23(1):316–326

2. Albarracín V, Kurth D, Ordoñez O, Belfiore C, Luccini E, Salum G, Piacentini R, Farías, ME. 2015. High-Up: A remote reservoir of Microbial Extremophiles in Central Andean Wetlands. Front Microbiol 6, 1404.

3. Dorador C, Vila I, Witzel KP, Imhoff JF. 2013. Bacterial and archaeal diversity in high altitude wetlands of the Chilean Altiplano. Fundam Appl Limnol 182, 135–159.

4. Aceituno P. 1997. Aspectos generales del clima en el Altiplano Sudamericano. In R Charrier, P Aceituno, M Castro, A Llanos, LA Raggi (Eds.), El Altiplano: ciencia y conciencia de los Andes, Actas del 21 simposio internacional de estudios altiplánicos (pp. 63–69). Santiago, Chile: Universidad de Chile.

5. Risacher F, Alonso H, Salazar C. 2003. The origin of brines and salts in Chilean salars: A hydrochemical review. Earth Sci Rev 63, 249–293.

6. Dorador C, Vila I, Imhoff JF, Witzel KP. 2008. Cyanobacterial diversity in Salar de Huasco, a high altitude saline wetland in northern Chile: an example of geographical dispersion?. FEMS microbiol 64(3), 419–432.

7. Hernández KL, Yannicelli B, Olsen LM, Dorador C, Menschel EJ, Molina, V, Remonsellez F, Hengst M, Jeffrey WH. 2016. Microbial activity response to solar radiation across contrasting environmental conditions in Salar de Huasco, Northern Chilean Altiplano. Front Microbiol 7, 1857.

8. Molina V, Hernández K, Dorador C, Eissler Y, Hengst M, Pérez V, Harrod C. 2016. Bacterial active community cycling in response to solar radiation and their influence on nutrient changes in a high-altitude wetland. Front Microbiol 7, 1–15.

9. Castro-Severyn J, Pardo-Esté C, Mendez KN, Morales N, Marquez SL, Molina F, Remonsellez F, Castro-Nallar E, Saavedra CP. 2020. Genomic Variation and Arsenic Tolerance Emerged as Niche Specific Adaptations by Different *Exiguobacterium* Strains Isolated From the Extreme Salar de Huasco Environment in Chilean– Altiplano. Front Microbiol 11, 1632.

10. Demergasso CS, Guillermo CD, Lorena EG, Mur JJP, Pedrós-Alió C. 2007. Microbial precipitation of arsenic sulfides in Andean salt flats. Geomicrobiol J 24, 111–123.

11. Escalante G, Campos V, Valenzuela C, Yañez J, Zaror C, Mondaca M. 2009. Arsenic resistant bacteria isolated from arsenic contaminated river in the Atacama Desert (Chile). Bull Environ Contam Toxicol 83(5):657H661.

12. Escudero LV, Casamayor EO, Chong G, Pedrós-Alió C, Demergasso C. 2013. Distribution of microbial arsenic reduction, oxidation and extrusion genes along a wide range of environmental arsenic concentrations. PLoS One 8(10), e78890.

13. Finstad KM, Probst AJ, Thomas BC, Andersen GL, Demergasso C, Echeverría A, Amundson R, Banfield JF. 2017. Microbial community structure and the persistence of cyanobacterial populations in salt crusts of the hyperarid Atacama Desert from genome-resolved metagenomics. Front Microbiol 8, 1435.

14. Wang L, Zhipeng Y, Jing C. 2020. Metagenomic insights into microbial arsenic metabolism in shallow groundwater of Datong basin, China. Chemosphere. 245. 125603.

15. Berhe A, Barnes R, Six J, Marin-Spiotta E. 2018. Role of soil erosion in biogeochemical cycling of essential elements: carbon, nitrogen, and phosphorus. Annu Rev Earth Pl Sci 46, 521–548.

16. Zhang J, Shi Q, Fan S, Zhang Y, Zhang M, Zhang J. 2020. Distinction between Cr and other heavy-metal-resistant bacteria involved in C/N cycling in contaminated soils of copper producing sites. J Hazard Mater 402, 123454.

17. Li Y, Zhang M, Xu R, Lin H, Sun X, Xu F, Gao P, Kong T, Xiao E, Yang N, Sun W. 2021. Arsenic and antimony co-contamination influences on soil microbial community composition and functions: Relevance to arsenic resistance and carbon, nitrogen, and sulfur cycling. Env Int 153, 106522.

18. Andres J, Bertin PN. 2016. The microbial genomics of arsenic. FEMS Microbiol Rev. 40:299–322.

19. Desoeuvre A, Casiot C, Héry M. 2016. Diversity and distribution of arsenic-related genes along a pollution gradient in a river affected by acid mine drainage. Microb Ecol 71(3), 672–685.

20. Rosen BP. 2002. Biochemistry of arsenic detoxification. FEBS Lett 529:86–92.

21. Sheik CS, Mitchell TW, Rizvi FZ, Rehman Y, Faisal M, Hasnain S, McInerney M, Krumholz, LR. 2012. Exposure of soil microbial communities to chromium and arsenic alters their diversity and structure. PloS one 7(6), e40059.

22. Kurth D, Amadio A, Ordoñez O, Albarracín V, Gärtner W, Farías ME. 2017. Arsenic metabolism in high altitude modern stromatolites revealed by metagenomic analysis. Sci Reports. 7: 1024.

23. Danczak RE, Johnston MD, Kenah C, Slattery M, Wilkins MJ. 2019. Capability for arsenic mobilization in groundwater is distributed across broad phylogenetic lineages. PloS one 14(9), e0221694.

24. Dunivin TK, Yeh SY, Shade A. 2019. A global survey of arsenic-related genes in soil microbiomes. BMC biology 17(1), 1–17.

25. Aykanat T, Lindqvist M, Pritchard VL, Primmer CR. 2016. From population genomics to conservation and management: A workflow for targeted analysis of markers identified using genome-wide approaches in Atlantic salmon Salmo salar. J Fish Biol 89(6), 2658–2679.

26. Ezeokoli OT, Bezuidenhout CC, Maboeta MS, Khasa DP, Adeleke RA. 2020. Structural and functional differentiation of bacterial communities in post-coal mining reclamation soils of South Africa: bioindicators of soil ecosystem restoration. Sci Rep 10(1), 1–14.

27. Dorador C, Vila I, Remonsellez F, Imhoff JF, Witzel KP. 2010. Unique clusters of Archaea in Salar de Huasco, an athalassohaline evaporitic basin of the Chilean Altiplano. FEMS Microbiol 73(2), 291–302.

28. Eissler Y, Dorador C, Kieft B, Molina V, Hengst M. 2020. Virus and Potential Host Microbes from Viral-Enriched Metagenomic Characterization in the High-Altitude Wetland, Salar de Huasco, Chile. Microorganisms 8(7), 1077.

29. Remonsellez F, Castro-Severyn J, Pardo-Esté C, Aguilar P, Fortt J, Salinas C, Barahona S, León J, Fuentes B, Areche C, Hernández K, Saavedra CP. 2018. Characterization and salt response in recurrent halotolerant *Exiguobacterium* sp. SH31 isolated from sediments of Salar de Huasco, Chilean Altiplano. Front Microbiol 9, 2228.

30. Vásquez-Dean J, Maza F, Morel I, Pulgar R, González M. 2020. Microbial communities from arid environments on a global scale. A systematic review. Biol Res 53(1), 1–12.

31. Cabrol NA, Grin EA, Chong G, Minkley E, Hock AN, Yu Y, Bebout L, Fleming E, Häder D, Demergasso C, Gibson J, Escudero L, Dorador C, Lim D, Woosley C, Morris R, Tambley T, Gaete V, Galvez M, Smith E, Uskin-Peate I, Salazar C, Dawidowicz G, Majerowicz J. 2009. The high-lakes project. J Geophys Res 114(G2).

32. Aguilar P, Acosta E, Dorador C, Sommaruga R. 2016. Large differences in bacterial community composition among three nearby extreme waterbodies of the high Andean plateau. Front Microbiol 7, 976.

33. Cortés-Albayay C, Silber J, Imhoff JF, Asenjo JA, Andrews B, Nouioui I, Dorador C. 2019. The polyextreme ecosystem, Salar de Huasco at the Chilean Altiplano of the Atacama Desert houses diverse Streptomyces spp. with promising pharmaceutical potentials. Diversity 11(5), 69.

34. Bourg ACM. 1989. Adsorption of Trace Inorganic and Organic Contaminants by Solid Particulate Matter, in Aquatic Ecotoxicology: Fundamental Concepts and Methodologies. Boudou A and Ribeyre F. eds. CRC Press, Inc. Boca Raton, Florida; pp.107–148.

35. Herrera V, De Gregori I, Pinochet H. 2009. Assessment of trace elements and mobility of arsenic and manganese in lagoon sediments of the salars of Huasco and Coposa, Chilean Altiplano. J Chil Chem Soc 54(3), 282–288.

36. Oren A. 2013. Life at high salt concentrations, intracellular KCl concentrations, and acidic proteomes. Front Microbiol 4, 315.

37. Farías ME, Rascovan N, Toneatti DM, Albarracín VH, Flores MR, Poiré DG, Collavino M, Aguilar M, Vazquez M, Polerecky L. 2013. The discovery of stromatolites developing at 3570 m above sea level in a high-altitude volcanic lake Socompa, Argentinean Andes. PloS one 8(1), e53497.

38. Farías ME, Contreras M, Rasuk MC, Kurth D, Flores MR, Poire DG, Novoa F, Visscher PT. 2014. Characterization of bacterial diversity associated with microbial mats, gypsum evaporites and carbonate microbialites in thalassic wetlands: Tebenquiche and La Brava, Salar de Atacama, Chile. Extremophiles 18(2), 311–329.

39. Saona LA, Soria M, Durán-Toro V, Wörmer L, Milucka J, Castro-Nallar E, Meneses C, Contreras M, Farías ME. 2021. Phosphate-Arsenic Interactions in Halophilic Microorganisms of the Microbial Mat from Laguna Tebenquiche: from the Microenvironment to the Genomes. Microb Ecol 1–13.

40. Ko MS, Park HS, Kim KW, Lee JU. 2013. The role of Acidithiobacillus ferrooxidans and Acidithiobacillus thiooxidans in arsenic bioleaching from soil. Environ Geochem HLTH 35(6), 727–733.

41. Demergasso C, Casamayor EO, Chong G, Galleguillos P, Escudero L, Pedros-Alió C. 2004 Distribution of prokaryotic genetic diversity in athalassohaline lakes of the Atacama Desert, Northern Chile. FEMS Microbiol Ecol 48:57–69

42. Demergasso C, Escudero L, Casamayor EO, Chong G, Balague V, Pedros-Alió C. 2008 Novelty and spatio-temporal heterogeneity in the bacterial diversity of hypersaline Lake Tebenquiche (Salar de Atacama). Extremophiles 12:491–504

43. Demergasso C, Dorador C, Meneses D, Blamey J, Cabrol N, Escudero L, Chong G. 2010 Prokaryotic diversity pattern in high-altitude ecosystems of the Chilean Altiplano. J Geophys Res 115:G00D09

44. Dorador C. 2007. Microbial communities in high altitude altiplanic wetlands in northern Chile: phytogeny, diversity and function. Doctoral dissertation, Christian-Albrechts-Universität, Kiel, Germany, p 166

45. Jiang H, Dong H, Yu B, Liu X, Li Y, Ji S, Zhang CL. 2007. Microbial response to salinity change in Lake Chaka, a hypersaline lake on Tibetan plateau. Environ Microbiol 9:2603–2621

46. Jung P, Baumann K, Lehnert LW, Samolov E, Achilles S, Schermer M, Wraase L, Eckhardt KU, Bader M, Leinweber P, Karsten U, Bendix J, Büdel B. 2020. Desert breath—How fog promotes a novel type of soil biocenosis, forming the coastal Atacama Desert’s living skin. Geobiology 18(1), 113–124.

47. Samolov E, Baumann K, Büdel B, Jung P, Leinweber P, Mikhailyuk T, Karsten U, Glaser K. 2020. Biodiversity of algae and cyanobacteria in biological soil crusts collected along a climatic gradient in Chile using an integrative approach. Microorganisms 8(7), 1047.

48. Castro-Severyn J, Remonsellez F, Valenzuela SL, Salinas C, Fortt J, Aguilar P, Pardo-Esté C, Dorador C, Quatrini R, Molina F, Aguayo D, Castro-Nallar E, Saavedra CP. 2017. Comparative genomics analysis of a new *Exiguobacterium* strain from Salar de Huasco reveals a repertoire of stress-related genes and arsenic resistance. Front Microbiol 8, 456.

49. Castro-Severyn J, Pardo-Esté C, Sulbaran Y, Cabezas C, Gariazzo V, Briones A, Morales N, Séveno M, Decourcelle, Salvetat N, Remonsellez F, Castro-Nallar E, Saavedra CP. 2019. Arsenic response of three altiplanic *Exiguobacterium* strains with different tolerance levels against the metalloid species: a proteomics study. Front Microbiol 10, 2161.

50. Idris H, Goodfellow M, Sanderson R, Asenjo JA, Bull AT. 2017. Actinobacterial rare biospheres and dark matter revealed in habitats of the Chilean Atacama Desert. Sci Rep 7(1), 1–11.

51. Agler MT, Ruhe J, Kroll S, Morhenn C, Kim ST, Weigel D, Kemen EM. 2016. Microbial hub taxa link host and abiotic factors to plant microbiome variation. PLoS biology 14(1), e1002352.

52. Kitahara R, Oyama K, Kawamura T, Mitsuhashi K, Kitazawa S, Yasunaga K, Sagara N, Fujimoto M, Terauchi, K. 2019. Pressure accelerates the circadian clock of cyanobacteria. Sci Rep 9(1), 1–8.

53. Crits-Christoph A, Gelsinger DR, Wierzchos J, Ravel J, Davila A, Casero MC, DiRuggiero J. 2016. Functional interactions of archaea, bacteria and viruses in a hypersaline endolithic community. Env Microbiol 18(6), 2064–2077.

54. Luton PE, Wayne JM, Sharp RJ, Riley PW. 2002. The *mcrA* gene as an alternative to 16S rRNA in the phylogenetic analysis of methanogen populations in landfill. Microbiology 148(11), 3521–3530.

55. Sela-Adler M, Ronen Z, Herut B, Antler G, Vigderovich H, Eckert W, Sivan O. 2017. Co-existence of methanogenesis and sulfate reduction with common substrates in sulfate-rich estuarine sediments. Front Microbiol 8, 766.

56. Xiao KQ, Li LG, Ma LP, Zhang SY, Bao P, Zhang T, Zhu YG. 2016. Metagenomic analysis revealed highly diverse microbial arsenic metabolism genes in paddy soils with low-arsenic contents. Env Poll 211, 1–8.

57. Zhai W, Qin T, Li L, Guo T, Yin X, Khan MI, Hashmi MZ, Liu X, Tang X, Xu, J. 2020. Abundance and diversity of microbial arsenic biotransformation genes in the sludge of full-scale anaerobic digesters from a municipal wastewater treatment plant. Env Int 138, 105535.

58. Slyemi D, Bonnefoy V. 2012. How prokaryotes deal with arsenic. Env Microbiol Rep 4(6), 571–586.

59. Jia Y, Huang H, Chen Z, Zhu YG. 2014. Arsenic uptake by rice is influenced by microbe-mediated arsenic redox changes in the rhizosphere. Env Sci Tech 48(2), 1001–1007.

60. Zhang SY, Zhao FJ, Sun GX, Su JQ, Yang XR, Li H, Zhu YG. 2015. Diversity and abundance of arsenic biotransformation genes in paddy soils from southern China. Env Sci Tech 49(7), 4138–4146.

61. Osman D, Cavet JS. 2010. Bacterial metal-sensing proteins exemplified by ArsR– SmtB family repressors. Nat Prod Rep 27(5), 668–680.

62. Shen Z, Luangtongkum T, Qiang Z, Jeon B, Wang L, Zhang Q. 2014. Identification of a novel membrane transporter mediating resistance to organic arsenic in *Campylobacter jejuni*. Antimicrob Agents Chemother 58(4), 2021–2029.

63. Fekih BI, Zhang C, Li YP, Zhao Y, Alwathnani HA, Saquib Q, Rensing C, Cervantes C. 2018. Distribution of Arsenic Resistance Genes in Prokaryotes. Front Microbiol 9, 2473.

64. Mateos LM, Villadangos AF, Alfonso G, Mourenza A, Marcos-Pascual L, Letek M, Pedre B, Messens J, Gil JA. 2017. The arsenic detoxification system in corynebacteria: basis and application for bioremediation and redox control. Adv Appl Microbiol 99, 103–137.

65. Shi K, Li C, Rensing C, Dai X, Fan X, Wang G. 2018. Efflux transporter ArsK is responsible for bacterial resistance to arsenite, antimonite, trivalent roxarsone, and methylarsenite. App Env Microbiol 84(24).

66. Xue S, Jiang X, Wu C, Hartley W, Qian Z, Luo X, Li W. 2020. Microbial driven iron reduction affects arsenic transformation and transportation in soil-rice system. Env Poll 260, 114010.

67. Oremland RS, Saltikov CW, Wolfe-Simon F, Stolz JF. 2009. Arsenic in the evolution of earth and extraterrestrial ecosystems. J Geomicrobiol 26(7), 522–536.

68. Mazumder P, Sharma SK, Taki K, Kalamdhad AS, Kumar M. 2020. Microbes involved in arsenic mobilization and respiration: a review on isolation, identification, isolates and implications. Environ Geochem HLTH 1–27.

69. Lobb B, Kurtz DA, Moreno-Hagelsieb G, Doxey AC. 2015. Remote homology and the functions of metagenomic dark matter. Front Genet 6, 234.

70. da Costa WLO, Araújo CLDA, Días LM, Pereira LCDS, Alves JTC, Araujo FA, Folador EL, Henriques I, Silva A, Folador ARC. 2018. Functional annotation of hypothetical proteins from the *Exiguobacterium antarcticum* strain B7 reveals proteins involved in adaptation to extreme environments, including high arsenic resistance. PloS one 13(6), e0198965.

71. Antczak M, Michaelis M, Wass MN. 2019. Environmental conditions shape the nature of a minimal bacterial genome. Nat comm 10(1), 1–13.

72. Makarova KS, Wolf YI, Koonin EV. 2019. Towards functional characterization of archaeal genomic dark matter. Biochem Soc Trans 47(1), 389–398.

73. Cai L, Liu G, Rensing C, Wang G. 2009. Genes involved in arsenic transformation and resistance associated with different levels of arsenic-contaminated soils. BMC Microbiol 9(1), 1–11.

74. Chen SC, Sun GX, Yan Y, Konstantinidis KT, Zhang SY, Deng Y, Li XM, Cui HL, Musat F, Popp D, Rosen B, Zhu YG. 2020. The Great Oxidation Event expanded the genetic repertoire of arsenic metabolism and cycling. PNAS 117(19), 10414–10421.

75. Delmont TO, Quince C, Shaiber A, Esen ÖC, Lee ST, Rappé MS, McLellan S, Lücker S, Eren AM. 2018. Nitrogen-fixing populations of Planctomycetes and Proteobacteria are abundant in surface ocean metagenomes. Nat Microbiol 3(7), 804–813.

76. Leema JM, Kirubagaran R, Vinithkumar NV, Dheenan PS, Karthikayulu S. 2010. High value pigment production from *Arthrospira* (Spirulina) platensis cultured in seawater. Bioresour Technol 101(23), 9221–9227.

77. Sorokin DY, Muntyan MS, Panteleeva AN, Muyzer G. 2012. *Thioalkalivibrio sulfidiphilus* sp. nov., a haloalkaliphilic, sulfur-oxidizing gammaproteobacterium from alkaline habitats. Int J Syst Evol Microbiol 62(8), 1884–1889.

78. Purcell AM, Mikucki JA, Achberger AM, Alekhina IA, Barbante C, Christner BC, Ghosh D, Michaud A, Mitchell A, Priscu J, Scherer R, Skidmore M, Vick-Majors TJ. 2014. Microbial sulfur transformations in sediments from Subglacial Lake Whillans. Front Microbiol 5, 594.

79. Zargar K, Hoeft S, Oremland R, Saltikov CW. 2010. Identification of a novel arsenite oxidase gene, *arxA*, in the haloalkaliphilic, arsenite-oxidizing bacterium *Alkalilimnicola ehrlichii* strain MLHE-1. J Bacteriol 192(14), 3755–3762.

80. Andrews S. 2010. FastQC a quality-control tool for high-throughput sequence data http://www.Bioinformaticsbabraham.ac.uk/projects/fastqc.

81. Bolger AM, Lohse M, Usadel B. 2014. Trimmomatic: a flexible trimmer for Illumina sequence data. Bioinformatics 30(15), 2114–2120.

82. Schmieder R, Edwards R. 2011. Quality control and preprocessing of metagenomic datasets. Bioinformatics 27(6), 863–864.

83. Gruber-Vodicka HR, Seah BK, Pruesse E. 2020. phyloFlash: Rapid Small-Subunit rRNA Profiling and Targeted Assembly from Metagenomes. Msystems, 5(5).

84. Quast C, Pruesse E, Yilmaz P, Gerken J, Schweer T, Yarza P, Peplies J, Glöckner FO. 2012. The SILVA ribosomal RNA gene database project: improved data processing and web-based tools. Nucleic Acids Res 41(D1), D590–D596.

85. R Core Team. 2018. R: A language and environment for statistical computing. R Foundation for Statistical Computing, Vienna, Austria. URL https://www.R-project.org/.

86. RStudio Team. 2016. RStudio: Integrated development for R. Boston, MA: RStudio Inc. Retrieved from http://www.rstudio.com/

87. Callahan B, McMurdie P, Rosen M, Han A, Johnson A, Holmes S. 2016. DADA2: High-resolution sample inference from Illumina amplicon data. Nat Methods 13, 581– 583.

88. Wang Q, Garrity GM, Tiedje JM, Cole JR. 2007. Naive Bayesian classifier for rapid assignment of rRNA sequences into the new bacterial taxonomy. Appl Environ Microbiol 73(16), 5261–5267.

89. Love M, Huber W, Anders S. 2014. Moderated estimation of fold change and dispersion for RNA-seq data with DESeq2. Genome Biol 15, 550.

90. Wright ES. 2016. Using DECIPHER v2. 0 to analyze big biological sequence data in R. R Journal 8(1).

91. Price MN, Dehal PS, Arkin AP. 2009. FastTree: computing large minimum evolution trees with profiles instead of a distance matrix. Mol Biol Evol 26(7), 1641–1650.

92. McMurdie PJ, Holmes S. 2013. phyloseq: an R package for reproducible interactive analysis and graphics of microbiome census data. PloS one 8(4), e61217.

93. Andersen KS, Kirkegaard RH, Karst SM, Albertsen M. 2018. ampvis2: an R package to analyse and visualise 16S rRNA amplicon data. BioRxiv 299537.

94. Wickham H. 2016. ggplot2: Elegant Graphics for Data Analysis. Springer-Verlag New York.

95. Lahti L, Shetty S, Blake T, Salojarvi J. 2017. Tools for microbiome analysis in R. Version, 1, 10013.

96. Gentleman R, Carey V, Huber W, Hahne F. 2011. Genefilter: Methods for filtering genes from microarray experiments. R package version, 1(0).

97. Kurtz ZD, Müller CL, Miraldi ER, Littman DR, Blaser MJ, Bonneau RA. 2015. Sparse and compositionally robust inference of microbial ecological networks. PLoS Comput Biol 11(5), e1004226.

98. Schloerke B, Crowley J, Cook D, Briatte F, Marbach M, Thoen E, Elberg A, Toomet O, Crowley J, Hofmann H, Wickman H. 2018. GGally: Extension to ‘ggplot2’. R package version 1.4.0. https://CRAN.R-project.org/package=GGally.

99. Overbeek R, Olson R, Pusch GD, Olsen GJ, Davis JJ, Disz T, Edwards R, Gerdes S, Parrello B, Shukla M, Vonstein V, Wattam A, Xia F, Stevens R. 2014. The SEED and the Rapid Annotation of microbial genomes using Subsystems Technology (RAST). Nucleic Acids Res 42(D1), D206–D214.

100. Silva GGZ, Green KT, Dutilh BE, Edwards RA. 2016. SUPER-FOCUS: a tool for agile functional analysis of shotgun metagenomic data. Bioinformatics 32(3), 354–361.

101. Buchfink B, Xie C, Huson DH. 2015. Fast and sensitive protein alignment using DIAMOND. Nat Methods 12(1), 59–60.

102. Parks DH, Tyson GW, Hugenholtz P, Beiko RG. 2014. STAMP: statistical analysis of taxonomic and functional profiles. Bioinformatics 30(21), 3123–3124.

103. Li D, Luo R, Liu CM, Leung CM, Ting HF, Sadakane K, Yamashita H, Lam TW. 2016. MEGAHIT v1.0: a fast and scalable metagenome assembler driven by advanced methodologies and community practices. Methods 102, 3–11.

104. Mikheenko A, Saveliev V, Gurevich A. 2016. MetaQUAST: evaluation of metagenome assemblies. Bioinformatics 32(7), 1088–1090.

105. Seemann T. 2014. Prokka: rapid prokaryotic genome annotation. Bioinformatics 30(14), 2068–2069.

106. Langmead B, Salzberg SL. 2012. Fast gapped-read alignment with Bowtie 2. Nat Methods 9(4), 357.

107. Li H, Handsaker B, Wysoker A, Fennell T, Ruan J, Homer N, Marth G, Abecasis G, Durbin R. 2009. The sequence alignment/map format and SAMtools. Bioinformatics 25(16), 2078–2079.

108. Eren AM, Esen ÖC, Quince C, Vineis JH, Morrison HG, Sogin ML, Delmont TO. 2015. Anvi’o: an advanced analysis and visualization platform for ‘omics data. PeerJ 3, e1319.

109. Hyatt D, Chen GL, LoCascio PF, Land ML, Larimer FW, Hauser LJ. 2010. Prodigal: prokaryotic gene recognition and translation initiation site identification. BMC Bioinformatics 11(1), 119.

110. Eddy SR. 2011. Accelerated profile HMM searches. PLoS Comput Biol 7(10), e1002195.

111. Tatusov RL, Galperin MY, Natale DA, Koonin EV. 2000. The COG database: a tool for genome-scale analysis of protein functions and evolution. Nucleic Acids Res 28(1), 33–36.

112. De Anda V, Zapata-Peñasco I, Poot-Hernandez AC, Eguiarte LE, Contreras-Moreira B, Souza V. 2017. MEBS, a software platform to evaluate large (meta) genomic collections according to their metabolic machinery: unraveling the sulfur cycle. GigaScience 6(11), gix096.

113. Anders S, Pyl PT, Huber W. 2015. HTSeq—a Python framework to work with high-throughput sequencing data. Bioinformatics 31(2), 166–169.

114. Aubry S, Kelly S, Kümpers BM, Smith-Unna RD, Hibberd JM. 2014. Deep evolutionary comparison of gene expression identifies parallel recruitment of trans-factors in two independent origins of C4 photosynthesis. PLoS Genet 10(6), e1004365.

115. Kolde R, Kolde MR. 2015. Package ‘pheatmap’. R Package 1(7), 790.

116. Katoh K, Misawa K, Kuma KI, Miyata T. 2002. MAFFT: a novel method for rapid multiple sequence alignment based on fast Fourier transform. Nucleic Acids Res 30(14), 3059–3066.

117. Alneberg J, Bjarnason BS, De Bruijn I, Schirmer M, Quick J, Ijaz UZ, Lahti L, Loman N, Andersson A, Quince C. 2014. Binning metagenomic contigs by coverage and composition. Nat Methods 11(11), 1144–1146.

118. Parks DH, Imelfort M, Skennerton CT, Hugenholtz P, Tyson GW. 2015. CheckM: assessing the quality of microbial genomes recovered from isolates, single cells, and metagenomes. Genome Res 25(7), 1043–1055.

119. Chaumeil PA, Mussig AJ, Hugenholtz P, Parks DH. 2020. GTDB-Tk: a toolkit to classify genomes with the Genome Taxonomy Database. Bioinformatics 36(6), btz848.

120. Parks DH, Chuvochina M, Chaumeil PA, Rinke C, Mussig AJ, Hugenholtz P. 2020. A complete domain-to-species taxonomy for Bacteria and Archaea. Nat Biotech 1–8.

121. Kanehisa M, Sato Y, Kawashima M, Furumichi M, Tanabe M. 2016. KEGG as a reference resource for gene and protein annotation. Nucleic Acids Res 44(D1), D457–D462.

122. Aramaki T, Blanc-Mathieu R, Endo H, Ohkubo K, Kanehisa M, Goto S, Ogata H. 2020. KofamKOALA: KEGG ortholog assignment based on profile HMM and adaptive score threshold. Bioinformatics 36(7), 2251–2252.

123. Muto A, Kotera M, Tokimatsu T, Nakagawa Z, Goto S, Kanehisa M. 2013. Modular architecture of metabolic pathways revealed by conserved sequences of reactions. J Chem Inf Model 53(3), 613–622.

124. Bastian M, Heymann S, Jacomy M. 2009. Gephi: an open-source software for exploring and manipulating networks. ICWSM 2, 361–362.

125. Bowers RM, Kyrpides NC, Stepanauskas R, Harmon-Smith M, Doud D, Reddy TBK, Schulz F, Jarett J, Rivers A, Eloe-Fadrosh EA, Tringe SG, Ivanova NN, Copeland A, Clum A, Becraft ED, Malmstrom RR, Birre B, Podar M, Bork P, Weinstock GM, Garrity G, Dodsworth JA, Yooseph S, Sutton G, Glöckner F, Gilbert J, Nelson W, Hallam S, Jungbluth SP, Ettema TJ, Tighe S, Konstantinidis KT, Liu WT, Baker BJ, Rattei T, Eisen JA, Hedlund B, McMahon KD, Fierer N, Knight R, Finn R, Cochrane G, Karsch-Mizrachi I, Tyson GW, Rinke C, Lapidus A, Meyer F, Yilmaz P, Parks D, Eren AM, Schriml L, Hugenholtz P, Woyke T. 2017. Minimum information about a single amplified genome (MISAG) and a metagenome-assembled genome (MIMAG) of bacteria and archaea. Nat Biotechnol 35(8), 725–731.

